# SARS-CoV-2 structural coverage map reveals state changes that disrupt host immunity

**DOI:** 10.1101/2020.07.16.207308

**Authors:** Seán I. O’Donoghue, Andrea Schafferhans, Neblina Sikta, Christian Stolte, Sandeep Kaur, Bosco K. Ho, Stuart Anderson, James Procter, Christian Dallago, Nicola Bordin, Matt Adcock, Burkhard Rost

## Abstract

In response to the COVID-19 pandemic, many life scientists are focused on SARS-CoV-2. To help them use available structural data, we systematically modeled all viral proteins using all related 3D structures, generating 872 models that provide detail not available elsewhere. To organise these models, we created a structural coverage map: a novel, one-stop visualization summarizing what is — and is not — known about the 3D structure of the viral proteome. The map highlights structural evidence for viral protein interactions, mimicry, and hijacking; it also helps researchers find 3D models of interest, which can then be mapped with UniProt, PredictProtein, or CATH features. The resulting Aquaria-COVID resource (https://aquaria.ws/covid) helps scientists understand molecular mechanisms underlying coronavirus infection. Based on insights gained using our resource, we propose mechanisms by which the virus may enter immune cells, sense the cell type, then switch focus from viral reproduction to disrupting host immune responses.

**Significance:** Currently, much of the COVID-19 viral proteome has unknown molecular structure. To improve this, we generated ∼1,000 structural models, designed to capture multiple states for each viral protein. To organise these models, we created a structure coverage map: a novel, one-stop visualization summarizing what is — and is not — known about viral protein structure. We used these data to create an online resource, designed to help COVID-19 researchers gain insight into the key molecular processes that drive infection. Based on insights gained using our resource, we speculate that the virus may sense the type of cells it infects and, within certain cells, it may switch from reproduction to disruption of the immune system.

## Introduction

Due to the COVID-19 pandemic, many life scientists have recently switched focus towards SARS-CoV-2 (Severe Acute Respiratory Syndrome Coronavirus 2). This includes structural biologists, who have so far determined ∼300 Protein Data Bank (PDB) entries (1) for the 27 viral proteins.

These structures are, in turn, driving molecular modeling studies, most focused on the spike glycoprotein (2). Some modelling studies focus on breadth of coverage, predicting 3D structures for the entire SARS-CoV-2 proteome; this has been done using AlphaFold (3), C-I-TASSER (4), Rosetta (5), and SwissModel (6). Unfortunately, these predictions vary greatly (7), raising accuracy concerns; additionally, these approaches derive few structural states for each viral protein, thereby producing moderate total numbers of models (e.g., 24 for AlphaFold and 116 for SwissModel).

We aim to address these limitations via a depth-based strategy that models multiple states for each viral protein, using all related 3D structures in the PDB — thus leveraging structures determined for other coronaviruses, such as SARS-CoV, BtCoV-HKU4 (bat coronavirus HKU4), FCoV (feline coronavirus), IBV (avian infectious bronchitis virus), and MHV-A59 (mouse hepatitis virus A59), as well as distant viruses, such as CHIKV (Chikungunya virus) and FMDV (foot-and-mouth disease virus).

Combining breadth and depth of coverage requires modeling methods with low computational cost; here, we use only sequence profile comparisons (8) to align viral sequences onto experimentally derived 3D structures (9). This generates what we call minimal models, in which 3D coordinates are not modified, but are mapped onto SARS-CoV-2 sequences using coloring that indicates model quality (10).

Minimal models have benefits: it is easy to understand how they were derived, helping assess the validity of insights gained. Thus, minimal models are broadly useful, even for researchers who are not modelling experts. Conversely, models generated by more sophisticated methods (3) can be more accurate, but require more time and expertise to assess their accuracy (7) and the validity of insights gained; this often limits their usefulness.

Large numbers of models can be generated by such minimal strategies, raising a new problem: how to visually organize such complex datasets to be usable. We consider this one instance of a critical issue impeding not just COVID-19 research, but all life sciences (11). Thus, we introduce a novel concept: a one-stop visualization summarizing what is known - and not known - about the 3D structure of the viral proteome. This tailored visualization — called the SARS-CoV-2 structural coverage map — helps researchers find structural models related to specific research questions.

Once a structural model of interest is found, it can be used to explore the spatial arrangement of sequence features — i.e., residue-based annotations, such as nonsynonymous mutations or post-translational modifications. Here, we integrated SARS-CoV-2 models into Aquaria (9), a molecular graphics system designed to simplify feature mapping and make minimal models broadly accessible to researchers who are not modelling experts. Previously, Aquaria could only map features from UniProt (12); for this work, we added >32,000 SARS-CoV-2 features from additional sources, and we also refactored Aquaria to improve performance.

The resulting Aquaria-COVID resource (https://aquaria.ws/covid) comprises a large set of SARS-CoV-2 structural information not readily available elsewhere. The resource also identifies structurally dark regions of the proteome, i.e., regions with no significant sequence similarity to any protein region observed by experimental structure determination (13). Clearly identifying such regions helps direct future research to reveal viral protein functions that are currently unknown.

Below, we describe the resource and systematically review insights it provides for each viral protein. We then discuss how these insights, taken together, reveal how viral proteins self-assemble, and how they may mimic (14) host proteins and hijack (15) host processes, including innate and adaptive immunity, nonsense-mediated decay, ubiquitination, and regulation of chromatin and telomeres.

During the COVID-19 pandemic, the Aquaria-COVID resource aims to fulfil a vital role by helping scientists more rapidly explore and assess evidence for the molecular mechanisms that underlie coronavirus infection — and to keep abreast of emerging knowledge, as new 3D structures and sequence features become available.

## Results

Our study was based on 14 UniProt (12) sequences that comprise the SARS-CoV-2 proteome (Table S1). We matched these against sequences of all available 3D structures in the PDB (1) — using profiles made with HHblits (8) — resulting in 872 minimal models (Tables S2-S4) that we incorporated into Aquaria (9), where they can be mapped with >32,000 features from UniProt, CATH (16), SNAP2 (17), and PredictProtein (18). These features include residue-based prediction scores for conservation, disorder, domains, flexibility, mutational propensity, subcellular location, and transmembrane helices (see Methods for details). To help other researchers use these models and features, we extensively refactored Aquaria to improve cross-platform performance, and created a matrix layout giving access to models for the 14 viral sequences (https://aquaria.ws/covid#matrix). We also created a structural coverage map (Figs. 1 & 2; https://aquaria.ws/covid) — a novel visual layout based on the viral genome organization. The coverage map summarizes key results obtained, including evidence for viral mimicry (Fig. 3) or hijacking of host proteins (Fig. 4), as well as viral protein interactions (Fig. 5). For each of the structural states shown in Fig. 2, a representative minimal model is specified via its Aquaria identifier in Table S5. In the following three sections of the Results, we systematically present key findings from structural models associated with three key regions of the viral genome; these sections are intended to be read with close reference to Fig. 2.

**Fig. 1.**
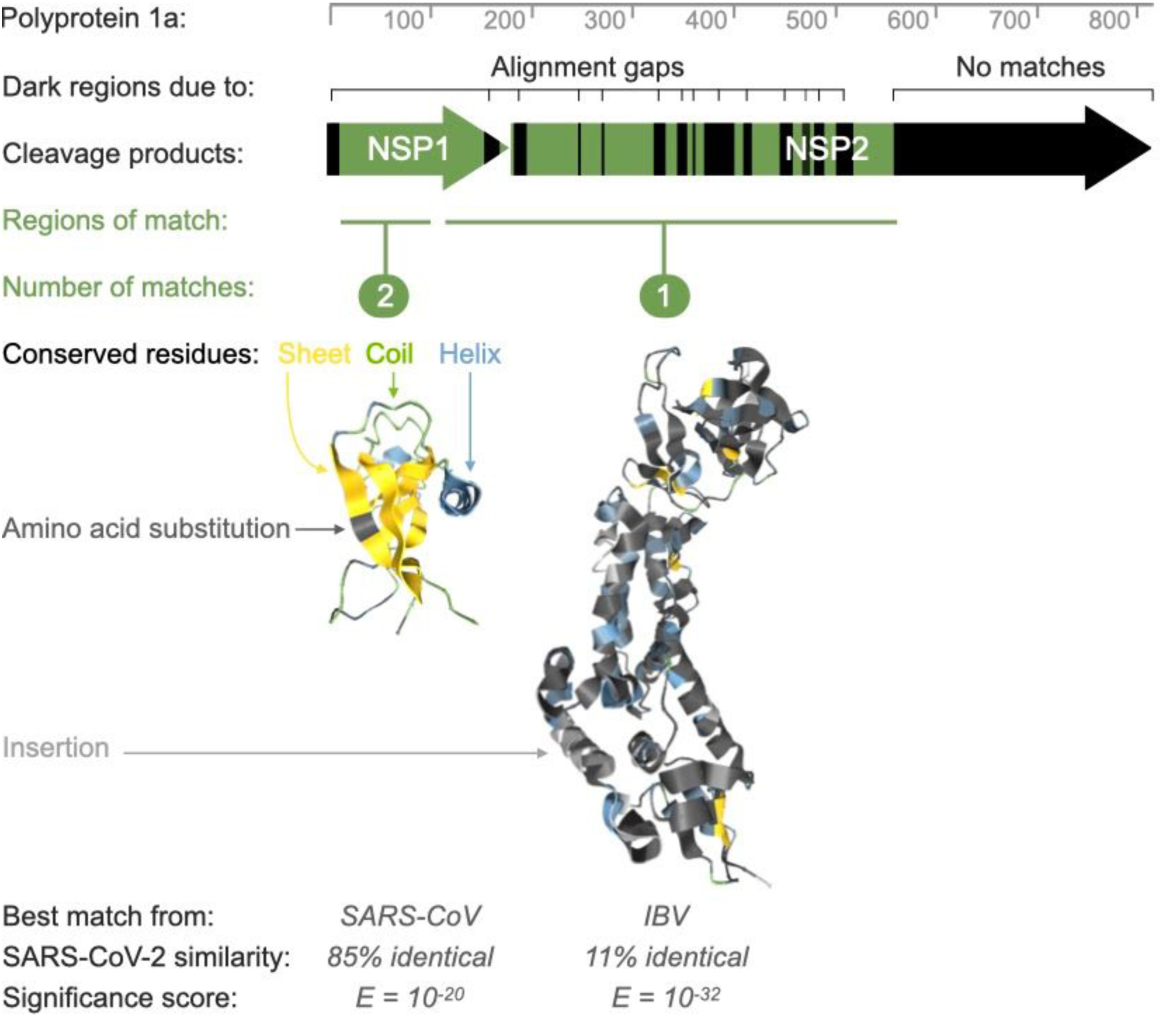
Graphical elements of the structural coverage map. The ruler (top) indicates sequence residue numbering. Dark sequence regions indicate either alignment gaps, or regions that do not match any experimentally-determined 3D structure. Regions with matching structure are indicated in green, and with a representative 3D structure position below, using dark gray coloring to indicate amino acid substitutions, using light gray to indicate alignment insertions, and using secondary structure coloring to indicate identical residues in the alignment. The coloring scheme makes it clear that the NSP1 model is similar in sequence to SARS-CoV-2, while the NSP2 model is remote. However, a better measure of model significance is the HHblits E-value, calculated by aligning sequence profiles for SARS-CoV-2 onto profiles for each 3D structure. The NSP2 model is more significant as it has a longer alignment. Made using Aquaria and edited with Keynote.

**Fig. 2.**
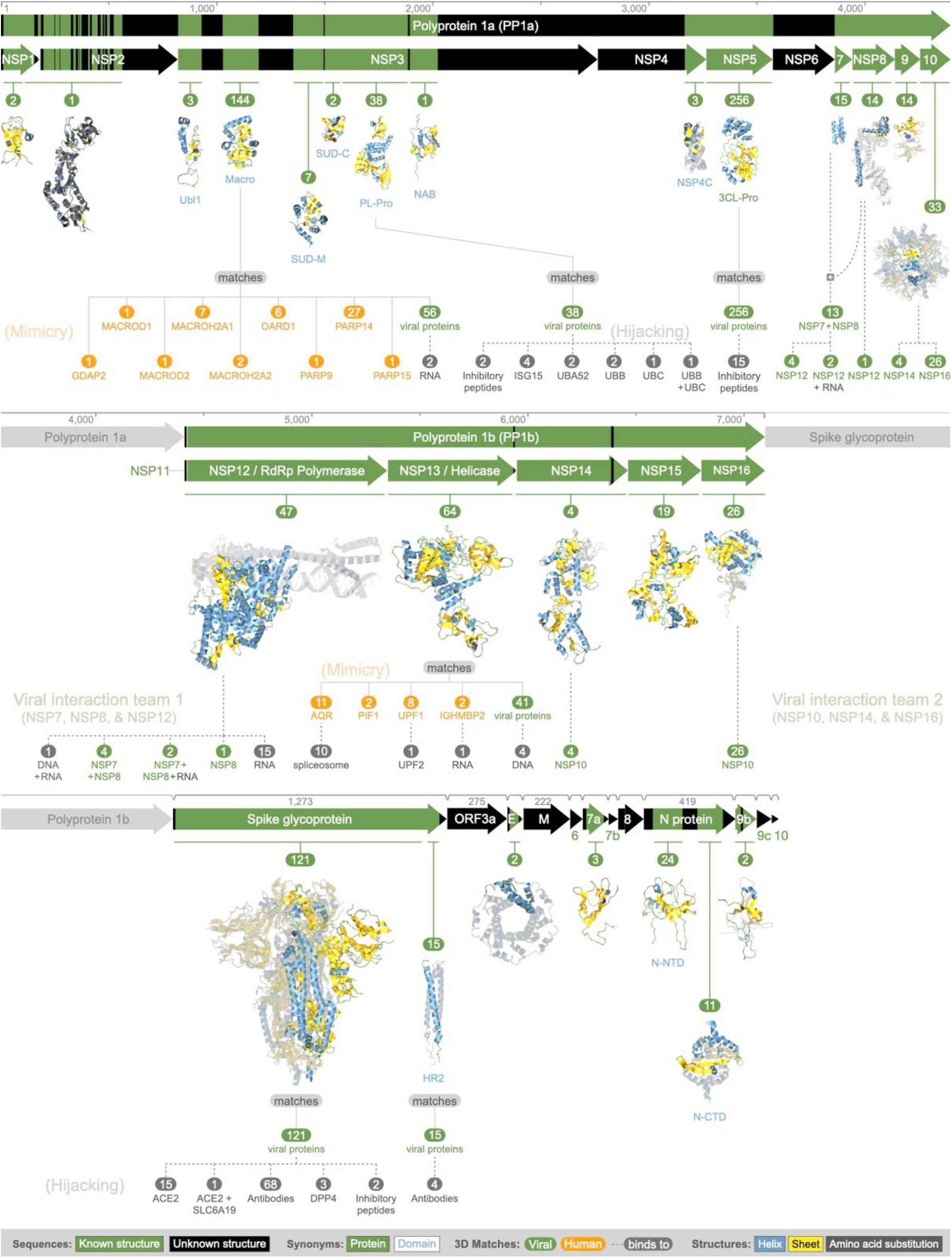
SARS-CoV-2 structural coverage map. Integrated visual summary of 3D structural evidence about viral protein states. Proteins are shown as arrows, scaled by sequence length, ordered by genomic location, and divided into three groups: (1) polyprotein 1a (top); (2) polyprotein 1b (middle); and (3) virion and accessory proteins (bottom). Dark coloring indicates sequence regions that do not match any experimentally-determined 3D structure. Regions with matching structure are indicated in green, and with representative 3D structures positioned below (Fig. 1), all at about the same physical scale. NSP3 (macro domain) and NSP13 may mimic the human proteins shown as orange nodes connected by solid lines (Fig. 3). NSP3 (PL-Pro domain) and spike glycoprotein may hijack the human proteins shown as gray nodes connected by dotted lines (Fig. 4). Interactions between viral proteins are shown using green colored nodes connected by dotted lines; these interactions occur in two disjoint teams: (1) NSP7, NSP8, and NSP12; and (2) NSP10, NSP14, and NSP16. For the remaining 18 viral proteins (‘suspects’), there was no structural evidence for interactions with other viral proteins, or for mimicry or hijacking of human proteins. Seven of the 18 are structurally dark proteins: NSP6, ORF3a, matrix glycoprotein, ORF6, ORF8, ORF9c, and ORF10. Details on the models found for each protein are presented in the Results, and in Table S5. Made using Aquaria and edited with Keynote.

**Fig. 3.**
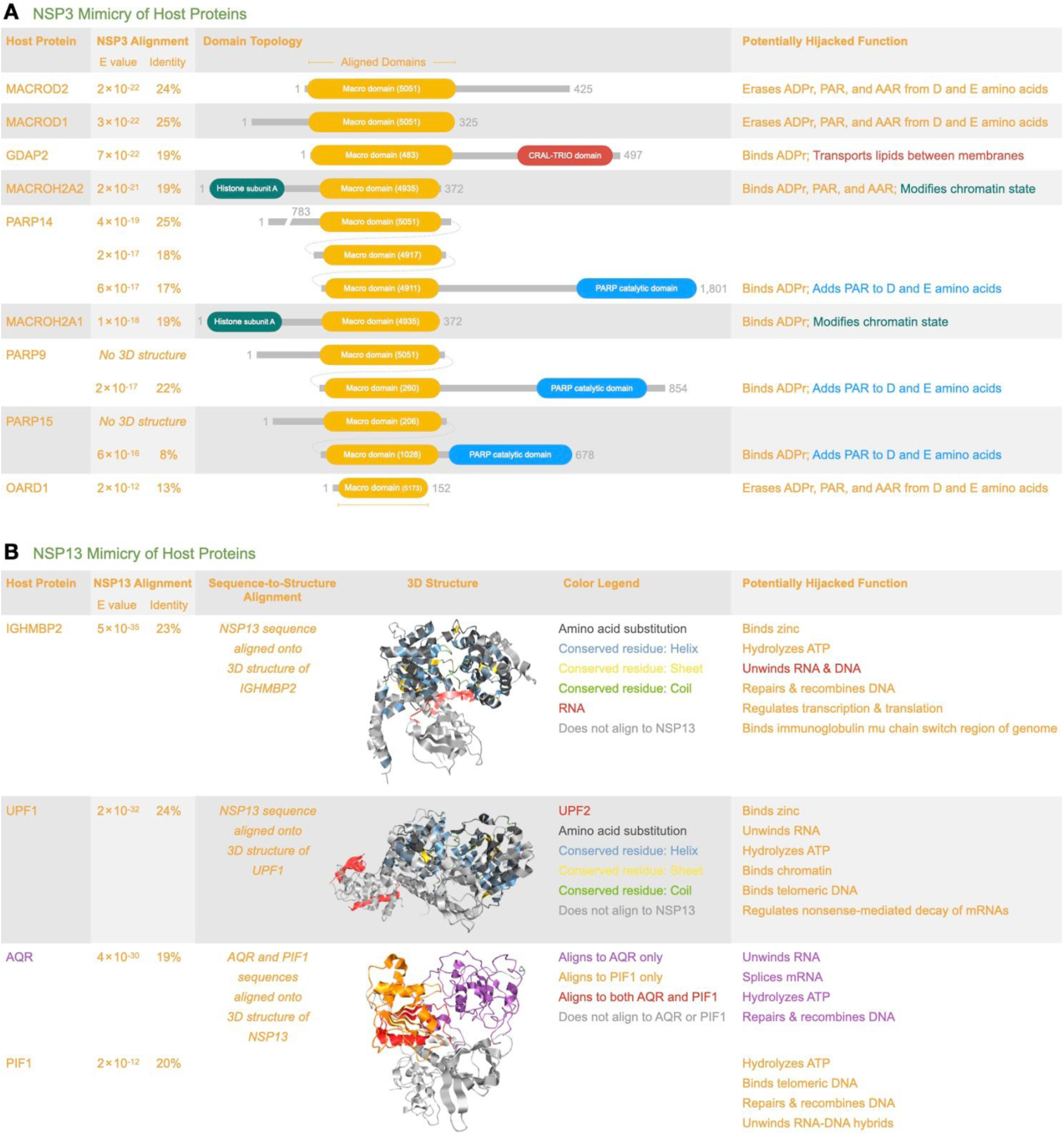
Viral mimicry of human proteins. (A) Lists domain topology for nine human proteins potentially mimicked by NSP3 (macro domain). The list was ranked by alignment significance, and includes a summary of potentially mimicked functions. Each macro domain is numbered to indicate its CATH functional family. The top ranked proteins (MACROD2 and MACROD1) remove ADPr from proteins, reversing the effect of ADPr writers (PARP14, PARP15, and PARP9), and affecting ADPr readers (GDAP2, MACROH2A2, and MACROH2A1). (B) Lists four human helicase proteins potentially mimicked by NSP13. The list was ranked by alignment significance, and includes a summary of potentially mimicked functions. There is stronger evidence (i.e., lower E value) for mimicry by NSP13 than by NSP3. For the two top ranked proteins, we show 3D structures of the human proteins, colored to indicate the region of alignment (Fig. 1) to NSP13. For IGHMBP2 (P0DTD1/4b3g), we see that RNA binds to the region matched by NSP13, suggesting that NSP13 binds RNA. For UPF1 (P0DTD1/2wjv), we see that UPF2 binds to a region not matched by NSP13, suggesting that NSP13 does not bind UPF2. For AQR and PIF1, we show a structure of NSP13 (P0DTD1/6jyt), colored to indicate regions matching AQR and PIF1. Made using features and structures from Aquaria, and edited in Photoshop and Keynote.

**Fig. 4.**
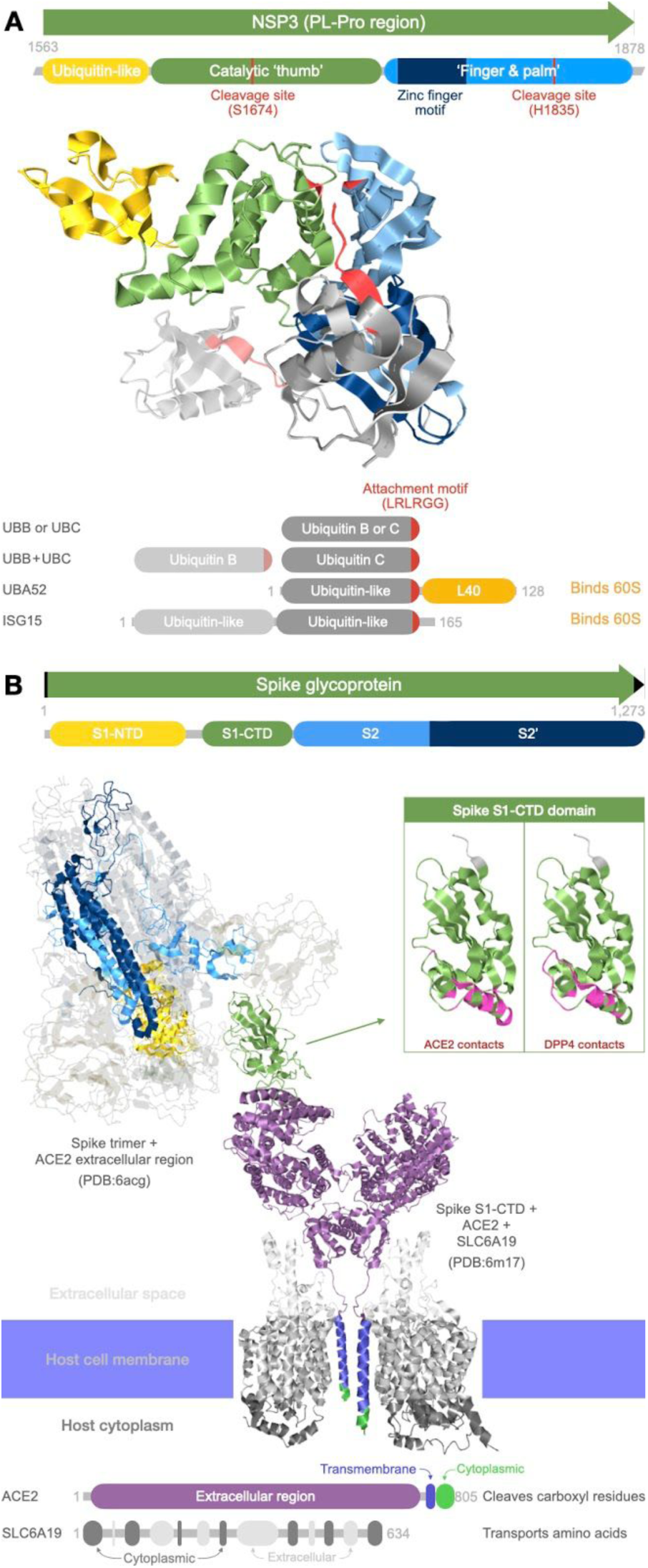
Viral hijacking of human proteins. (A) NSP3 (PL-Pro region) has three domains (yellow, green, and blue) that form two binding sites for ubiquitin-like (Ubl) domains (gray). In most matching structures, a single Ubl domain occupies the primary binding site (dark gray), with its C-terminal LRLRGG motif (red) positioned between the PL-Pro thumb and palm domains, and ending at the proteolytic cleavage site (red). The LRLRGG motif can attach to other proteins, including other copies of ubiquitin. PL-Pro cleaves this attachment, reversing mono- or poly-ubiquitination. One matching structure showed binding of two ubiquitin domains (UBB and UBC), with UBB occupying the secondary site (light gray). Other structures showed binding to the N- and C-terminal Ubl domains of ISG15 (interferon stimulated gene 15) at the secondary and primary sites, respectively. (B) The spike trimer structure (top left; P0DTC2/6acg) shows the pre-fusion state after cleavage at the S1/S2 boundary, and with one S1-CTD domain (green) bound to the extracellular region of the carboxypeptidase ACE2 (purple). Below that, we have manually aligned a second structure (P0DTC2/6m17) showing S1-CTD in complex with ACE2 and the amino acid transporter SLC6A19. By integrating sequence features (top and bottom) with 3D structures, the resulting figure shows the cellular context of the pre-fusion event, when spike first binds the ACE + SLC6A19 complex. The insert highlights S1-CTD residues that contact ACE2 (magenta). These are compared to residues that contact the N-terminal dipeptide peptidase DPP4 in a structure from BtCoV-HKU4 (P0DTC2/4qzv). DPP4 may also be targeted by SARS-CoV-2 (76); if so, our analysis shows that the S1-CTD + DPP4 binding site may partly overlap the S1-CTD + ACE2 binding site. Made using features and structures from Aquaria, and edited in Photoshop and Keynote.

**Fig. 5.**
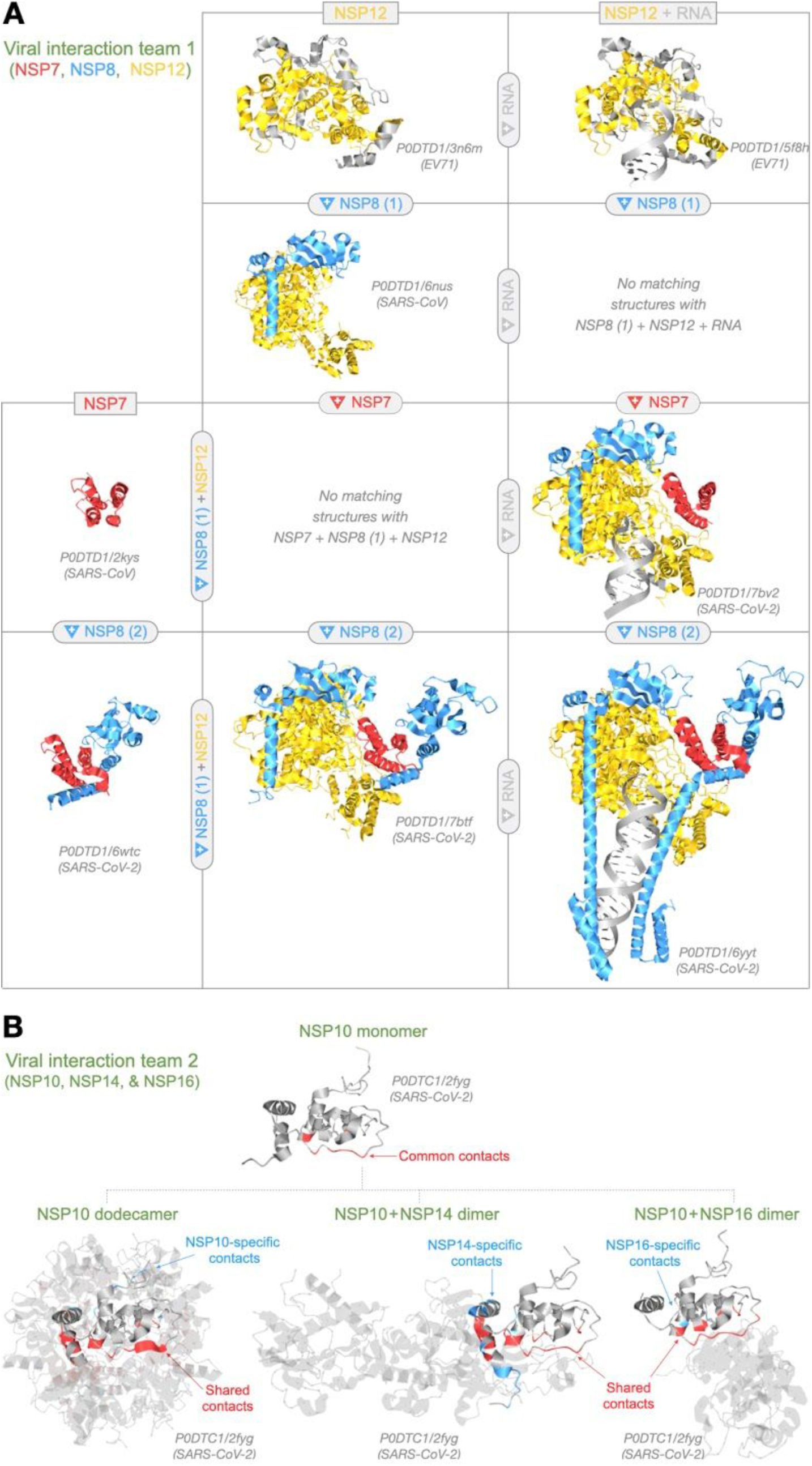
Viral protein interactions. (A) Summarizes all structural states observed for NSP7 (red), NSP8 (blue), and NSP12 (yellow). NSP12 alone (top row, left) can replicate RNA (top row, right); however, all matching structures were from distantly related viruses (pairwise identity ≤ 18%), such as EV71 (enterovirus). NSP8 binds NSP12 at two sites: (1) at the NSP12 core (2nd row, left); and (2) via NSP7-mediated cooperative interactions at the NSP12 periphery (bottom row, center), greatly enhancing RNA replication (bottom row, right). By themselves, NSP7 + NSP8 (bottom row, left) can also replicate RNA (25). Six of the matching structures showed NSP7 + NSP8 as a dimer; however, one structure showed a tetramer and one other showed a hexadecamer (P0DTD1/3ub0 and P0DTD1/2ahm, respectively). (B) Summarizes all structural states observed for NSP10, NSP14, and NSP16 (Team 2). One matching structure showed an NSP10 monomer (top), while two structures showed NSP10 as a dodecamer (bottom, left). All structures matching NSP14 showed a heterodimer with NSP10 (bottom, middle). All structures matching NSP16 showed a heterodimer with NSP10 (bottom, right). Nine NSP10 residues (shown in red on the monomer) were common intermolecular contacts in all three oligomers. Each oligomer also had specific NSP10 contacts (blue); but most NSP10 contacts were shared with at least one other oligomer (shown in red on each oligomer). This suggests that NSP10, NSP14, and NSP16 interact competitively. Made using Aquaria and edited with Keynote.

### Polyprotein 1a

Polyprotein 1a (a.k.a. PP1a) derives from polyprotein 1ab (a.k.a. PP1ab), and is cleaved into 10 proteins (NSP1–NSP10) that modify viral proteins, disable host defenses, and support viral replication.

NSP1 derives from residues 1–180 of PP1a and is thought to interact with the ribosome, suppressing translation of host mRNAs and promoting their degradation (19). We found two structures matching most of the NSP1 sequence (Fig. 1), both determined for NSP1 from SARS-CoV, which had 85% pairwise sequence identity to the matched region of NSP1 from SARS-CoV-2. These sequence-to-structure alignments were generated using HHblits sequence profiles (8), and had a significance score (E-value) of 10^−20^, giving an estimated ≥ 97% likelihood that NSP1 from SARS-CoV and SARS-CoV-2 adopt similar structures (9). The more recent structure (with Aquaria identifier P0DTC1/2gdt) was selected for the representative image (Figs. 1 & 2), which was colored to convey alignment quality (10). Unfortunately, neither structure showed interactions with other proteins or RNA and — unusually — they provided few functional insights (20), partly because they had a unique fold with no matches in CATH (16). NSP1 also had two short dark regions resulting from alignment gaps (Figs. 1 & 2); the N-terminal dark region may be accounted for by high flexibility (average predicted B-value = 60; see Methods).

NSP2 (PP1a 181–818) may disrupt intracellular signaling by interacting with host proteins (21). Unfortunately, these interactions are not captured in the single matching structure (Figs. 1 & 2) derived for NSP2 from IBV, a remote homolog with 11% pairwise sequence identity to the matched region of PP1a (residues 104–562). However, the HHblits E-value was 10^−32^, giving ≥ 98.5% likelihood that SARS-CoV-2 NSP2 adopts a similar structure — a stronger prediction than for NSP1 (above), largely because the matched region was much longer. Interestingly, the matched region partly overlapped NSP1 (PP1a 104–180). NSP2 also had multiple short dark regions, resulting from alignment gaps, plus a long C-terminal dark region (Figs. 1 & 2). The Aquaria webpage for this matching structure showed that — unusually — there is no scientific publication associated with this 10 year-old dataset (Table S5). Thus, while it may be of some use for researchers focused on NSP2, for most researchers this structure should probably be excluded, making NSP2 an entirely dark protein.

NSP3 (PP1a 819–2763) is a large, multidomain protein thought to perform many functions, including anchoring the viral replication complex to double-membrane vesicles derived from the endoplasmic reticulum (22). NSP3 had 188 matching structures clustered in nine distinct sequence regions.

NSP3 region 1 (a.k.a. Ubl1; PP1a 819–929) was the least conserved NSP3 region (average ConSurf score = 3.7; see Methods), suggesting it adapts to host-specific defenses. Ubl1 is thought to bind single-stranded RNA and the viral nucleocapsid protein (22). Unfortunately, these interactions are not shown in the three matching structures found, which all adopt a ubiquitin-like topology (CATH 3.10.20.350). Although it has distinct structural differences, Ubl1 may mimic host ubiquitin (22); however, we found no matches to structures of human ubiquitin, undermining the mimicry hypothesis. For the representative image (Fig. 2), we used the top-ranked matching structure provided in the Aquaria interface; further details about this and all models mentioned below are given in Table S5.

NSP3 region 2 (PP1a 930–1026) had no matches in CATH and no matching structures. This was the NSP3 region with lowest predicted sensitivity to mutation (median sensitivity 0%; see Methods), highest predicted flexibility (average B-factor = 55), highest fraction of disordered residues (47%), and highest fraction of residues predicted to be solvent-accessible (99%). We speculate that this region acts as a flexible linker and may contain post-translational modification sites hijacking host signalling, as are often found in viral disordered regions (15).

NSP3 region 3 (PP1a 1027–1193) has a macro domain (CATH 3.40.220.10) that may counteract innate immunity via interfering with ADP-ribose (ADPr) modification (22). This was the second least conserved NSP3 region (ConSurf = 3.9) and had the highest fraction of mutationally sensitive residues (29%), suggesting it is well adapted to specific hosts. This region had 144 matching structures (Fig. 2; Table S5). Of the matching structures, 47 were of human proteins (Figs. 2 & 3A), suggesting these host proteins may be mimicked by the virus. The potentially mimicked human proteins were: GDAP2, MACROD1, MACROD2, MACROH2A1, MACROH2A2, OARD1, PARP9, PARP14, and PARP15. An additional 56 structures matched to viral proteins, two in complex with RNA. For brevity here and in Fig. 2, we have omitted the remaining 41 matching structures derived from other host organisms (Table S2).

NSP3 region 4 (PP1a 1027–1368) comprised a disordered region plus a macro-like domain called SUD-N (CATH 3.40.220.30) that may bind RNA (22). This region had no matching structures.

NSP3 region 5 (PP1a 1389–1493) comprised another macro-like domain called SUD-M (CATH 3.40.220.20) that may bind both RNA and host proteins, and take part in viral replication (22). However, no interactions are shown in the seven matching structures found. Comparing these with structures matching NSP3 region 3, we see considerable differences and no evidence of mimicry of host macro domains (Fig. 2).

NSP3 region 6 (PP1a 1494–1562) comprised a domain called SUD-C (CATH 2.30.30.590) and had two matching structures, one of which also spanned NSP3 region 7. Using these structures, Lei et al. speculated that this region may bind metal ions and induce oxidative stress (22).

NSP3 region 7 (PP1a 1563–1878) comprises a papain-like protease (a.k.a. PL-Pro), made up of three domains (CATH 3.10.20.540, 1.10.8.1190, and 3.90.70.90) thought to cleave NSP1–NSP3 from the polyprotein and to cleave ubiquitin-like modifications from host proteins (Fig. 4A), thereby undermining interferon-induced antiviral activity (22). PL-Pro had 38 matching structures, of which 11 show binding to human ubiquitin-like proteins (Figs. 2 & 4A): four showed binding to ISG15; two showed binding to UBA52; two showed binding to UBB; one showed binding to UBC, and one showed binding to both UBB and UBC. Two additional matching structures showed the PL-Pro region in complex with inhibitory peptides.

NSP3 region 8 (PP1a 1879–2020) comprises a nucleic-acid binding domain (a.k.a. NAB) thought to bind single-stranded RNA and to unwind double-stranded DNA (22). NAB had only one matching structure with a fold not seen in any other PDB structure (CATH 3.40.50.11020).

NSP3 region 9 (PP1a 2021–2763) may anchor NSP3 to double-membrane vesicles (22). This was the most conserved NSP3 region (ConSurf = 4.9), suggesting it is less adapted to specific hosts. This region had no CATH matches and no matching structures.

NSP4 (PP1a 2764–3263) may act with NSP3 and NSP6 to create the double-membrane vesicles required for viral replication (23). NSP4 mostly comprised a dark region (PP1a 2764–3172) with no CATH matches, no disorder, and multiple transmembrane helices. The C-terminal region (PP1a 3173–3263) comprised a domain called NSP4C (CATH 1.10.150.420) that had three matching structures; all were homodimers, although NSP4C is thought to act as a monomer (24).

NSP5 (a.k.a. 3CL-Pro; PP1a 3264–3569) is a two-domain protein (CATH 2.40.10.10 and 1.10.1840.10) thought to cleave the viral polyprotein at 11 sites, resulting in NSP5–NSP16. NSP5 had 256 matching structures — more than any other viral protein — of which, 15 showed binding to inhibitory peptides.

NSP6 (PP1a 3570–3859) is a transmembrane protein thought to act with NSP3 and NSP4 to create double-membrane vesicles (23). Of the 15 PP1ab proteins, NSP6 was the only completely dark protein and was the least conserved (ConSurf = 2.9). NSP6 also had no CATH matches and no disordered regions.

NSP7 (PP1a 3860–3942) may support viral genome replication (25); it had 15 matching structures, some showing interactions with other viral proteins (Figs. 2 & 5A). In two structures, NSP7 occurred as a monomer, while the remaining 13 structures showed NSP7 bound to NSP8. Six of these structures also showed binding to NSP12; two of these six structures also showed binding to RNA. NSP7 comprises an antiparallel helical bundle (CATH 1.10.8.370) that adopts distinct substates, depending on its interaction partners (Fig. 5A).

NSP8 (PP1a 3943–4140) is also thought to support viral genome replication (25). It features a highly conserved (ConSurf = 7.3) ‘tail’ segment (PP1a 3943–4041), predominantly helical with some disordered residues and no CATH matches, followed by a less conserved (ConSurf = 5.7) ‘head’ domain (PP1a 4042–4140), with a beta barrel fold (CATH 2.40.10.290). NSP8 had 14 matching structures, all showing interactions to other viral proteins (Figs. 2 & 5A). One structure showed binding to NSP12 only, while the remaining 13 structures all showed binding to NSP7. Six of these 13 structures also showed binding to NSP12, and two of these six showed binding to RNA as well.

NSP9 (PP1a 4141–4253) may bind single-stranded RNA and take part in genome replication (26). NSP9 had 14 matching structures, all with beta barrel architecture (CATH 2.40.10.250) and mostly homodimers — thought to be the functional state (26).

NSP10 (PP1a 4254–4392) is thought to act with NSP14 and NSP16 to cap and proofread RNA during genome replication (27). NSP10 had no CATH matches, yet had 33 matching structures, some showing interactions to other viral proteins (Figs. 2 & 5B). In one matching structure, NSP10 was monomeric, while in two structures, NSP10 was a homododecamer that formed a hollow sphere (Fig. 1). Four matching structures showed binding to NSP14, while the remaining 26 structures showed binding to NSP16.

### Polyprotein 1b

Polyprotein 1b (a.k.a. PP1b) is cleaved by 3CL-Pro into five proteins (NSP12–NSP16) that drive replication of viral RNA. These proteins were predicted to have no disordered regions, no transmembrane helices, very few dark regions, and to be highly conserved (ConSurf = 5.3–6.6), compared with PP1a proteins (ConSurf = 2.9–5.6).

NSP12 (PP1ab 4393–5324) is an RNA-directed RNA polymerase (a.k.a. RdRp) thought to drive viral genome replication (28). NSP12 was the second most conserved PP1ab protein (ConSurf = 6.5), and had 47 matching structures, many showing interactions to other viral proteins (Figs. 2 & 5A). One structure showed binding to NSP8, while four structures showed binding to both NSP8 and NSP7; two of these four also showed binding to RNA. An additional 15 structures showed binding to RNA only, while one structure showed binding to both RNA and DNA.

NSP13 (PP1ab 5325–5925) is a multi-functional helicase thought to unwind double-stranded RNA and DNA, and to be a core component of the viral replication complex (29). The N-terminal half of NSP13 (PP1ab 5325–5577) had no matches in CATH, while the C-terminal half contained two Rossman fold domains (CATH 3.40.50.300). NSP13 had 64 matching structures, of which 23 showed potential mimicry of four human helicase proteins (Figs. 2 & 3B). Two of these structures showed matching between PIF1 and the first Rossman fold domain (PP1ab 5597–5733). A further 11 structures showed matching between AQR and the second Rossman fold domain plus part of the first; of these structures, 10 showed AQR bound to the spliceosome. The remaining 10 structures matched to both Rossman fold domains: eight of these structures showed mimicry of UPF1, of which two structures also showed binding to UPF2; finally, two structures showed mimicry of IGHMBP2, of which one structure also showed binding with RNA. A further 41 structures showed matches to viral proteins, of which four structures also showed binding to DNA.

NSP14 (PP1ab 5926–6452) is a proofreading exoribonuclease thought to remove 3’-terminal nucleotides from RNA, thereby reducing mutations during viral genome replication (30). NSP14 had no matches in CATH, but had four matching structures, all in complex with NSP10 (Figs. 2 & 5B).

NSP15 (PP1ab 6453–6798) is a uridylate-specific endoribonuclease thought to support viral genome replication (31). The N-terminal region of NSP15 had two domains (CATH 2.20.25.360 and 3.40.50.11580), while the C-terminal region (PP1ab 6642–6798) had none. NSP15 had 19 matching structures, none showing matching to — or interactions with — human proteins, or with other viral proteins (Fig. 2).

NSP16 (PP1ab 6799–7096) may methylate viral mRNA caps, following replication, which is thought to be important for evading host immune defenses (32). This was also the most conserved PP1ab protein (ConSurf = 6.6). NSP16 had a Rossman fold domain (CATH 3.40.50.150) that had 26 matching structures, all showing binding to NSP10 (Figs. 2 & 5B).

### Virion and Accessory Proteins

The 3’ end of the genome encodes 12 proteins, many involved in virion assembly. Remarkably, our analysis found no interactions between these proteins.

Spike glycoprotein binds host receptors, thereby initiating membrane fusion and viral entry (33). This protein has four UniProt domains, of which one (S1-CTD; residues 334–527) binds to host receptors (Fig. 4B). Aquaria found 136 matching structures clustered in two regions (Fig. 2). One was a heptad repeat region (HR2), with 15 matching structures, of which four structures showed binding to antibodies. The second region had 121 matching structures, of which 68 showed binding to antibodies and two structures showed binding to inhibitory peptides. Of the remaining structures, 18 showed interaction with human proteins (Figs. 2 & 4B): 15 showed binding to ACE2, of which one structure also showed binding to SLC6A19; finally, three structures showed binding to DPP4.

ORF3a may act as a homotetramer, forming an ion channel in host cell membranes that helps with virion release (34). ORF3a had no matching structures.

The envelope protein (a.k.a. E protein) matched two structures; one was a monomer while the other was a pentamer, forming an ion channel thought to span the viral envelope (35).

The matrix glycoprotein (a.k.a. M protein) is also thought to be part of the viral envelope (36). The matrix glycoprotein had no matching structures.

ORF6 may block expression of interferon stimulated genes (e.g., ISG15) that have antiviral activity (37). ORF6 had no matching structures.

ORF7a may interfere with the host cell surface protein BST2, preventing it from tethering virions (38). ORF7a had three matching structures.

ORF7b is an integral membrane protein thought to localize to the Golgi compartment and the virion envelope (39). ORF7b had no matching structures.

ORF8 is thought to inhibit type 1 interferon signaling (40); it is also very different to proteins from other coronaviruses. ORF8 had no matching structures.

The nucleocapsid protein (a.k.a. N protein) is thought to package the viral genome during virion assembly through interaction with the matrix glycoprotein, and also to become ADP-ribosylated (41). Depending on its phosphorylation state, this protein may also switch function, translocating to the nucleus and interacting with the host genome (42). This protein had 35 matching structures clustered in two regions: the N-terminal region had 24 matching structures, of which one was a tetramer, five were dimers, and the rest were monomers; the C-terminal region had 13 matching structures, all dimers.

ORF9b is a lipid-binding protein thought to interact with mitochondrial proteins, thereby suppressing interferon-driven innate immunity responses (43). ORF9b matched two structures that showed binding to a lipid analog (44).

ORF9c (a.k.a. ORF14) is currently uncharacterized experimentally; it is predicted to have a single-pass transmembrane helix. ORF9c had no matching structures.

ORF10 is a predicted protein that currently has limited evidence of translation (45), has no reported similarity to other coronavirus proteins, and has no matching structures.

## Discussion

The 872 models derived in this study capture essentially all structural states of SARS-CoV-2 proteins with supporting experimental evidence. We used these states to create a structural coverage map (Fig. 2): a concise yet comprehensive visual summary of what is known about the 3D structure of the viral proteome. Remarkably, we found so few states showing viral self-assembly (Fig. 5), mimicry (Fig. 3), or hijacking (Fig. 4) that — excluding non-human host proteins — all states could be included in the coverage map via several simple graphs. This may indicate that host interactions are rarely used in COVID-19 infection, consistent with the notion that viral activity is largely shielded from the host. However, other experimental techniques have found many more interactions between viral proteins (46), and with host proteins (45). Thus, the small number of interactions found in this work likely indicates limitations in current structural data.

Using the current structural data (Fig. 2), we can divide the 27 SARS-CoV-2 proteins into four categories: mimics, hijackers, teams, and suspects — below, we highlight insights derived in this work for each of these categories.

### Mimics

We found structural evidence for mimicry of human proteins for only two SARS-CoV-2 proteins: NSP3 and NSP13 (Figs. 2 & 3; Table S6).

NSP3 may mimic host proteins containing macro domains, interfering with ADP-ribose (ADPr) modification and thereby suppressing host innate immunity (22). We found nine potentially mimicked proteins (Fig. 3A); the top ranked matches (MACROD2 and MACROD1) remove ADPr from proteins (47), reversing the effect of ADPr writers (e.g., PARP9, PARP14, and PARP15 in lymphoid tissues), and affecting ADPr readers (e.g., the core histone proteins MACROH2A1, and MACROH2A2, found in most cells). Thus we speculate that, in infected cells, ADPr erasure by NSP3 may influence epigenetic regulation of chromatin state (48), potentially contributing to variation in COVID-19 patient outcomes. Furthermore, in infected macrophages, activation by PARP9 and PARP14 may be undermined by NSP3’s erasure of ADPr, resulting in vascular disorders (49), as seen in COVID-19 (50).

NSP13 may mimic four human helicases, based on stronger alignment evidence than for mimicry by NSP3 (Fig. 3). Three of these helicases are associated with DNA repair and recombination, thus we speculate mimicry of these proteins could either recruit host proteins to assist in viral recombination (51), or alternatively, could dysregulate recombination of host DNA. However, we found no evidence for mimicry of the ∼100 other human helicases (52), many also involved in recombination, suggesting that mimicry may hijack more specific functions performed by the four helicases. The top ranked helicase was IGHMBP2 (a.k.a. SMBP2), which acts in the cytoplasm as well as the nucleus, where it interacts with single-stranded DNA in the class switching region of the genome (53), close to *IGMH*, the gene coding the constant region of immunoglobulin heavy chains. We speculate that mimicry of IGHMBP2 may explain the dysregulation of immunoglobulin-class switching observed clinically (54). The second ranked helicase was UPF1 (a.k.a. regulator of nonsense transcripts 1, or RENT1), which acts in the cytoplasm as part of the nonsense-mediated mRNA decay pathway, known to counteract coronavirus infection (55); we speculate that mimicry of UPF1 may hijack this pathway, thus impeding host defenses. UPF1 also acts in the nucleus, interacting with telomeres — as does the helicase PIF1; we speculate that mimicry of UPF1 or PIF1 may be implicated to the connection seen between COVID-19 severity, age, and telomere length (56). In summary, we propose that (as mentioned above for nucleocapsid protein) NSP13 may sometimes switch from its key role in viral replication to a state focused on host genome interactions that undermine host immunity.

### Hijackers

We found direct structural evidence for hijacking of human proteins for only two SARS-CoV-2 proteins: NSP3 and spike glycoprotein (Figs. 2 & 4; Table S6).

NSP3 is believed to cleave ubiquitin from host proteins, thereby suppressing innate immune response and disrupting proteasome-mediated degradation (22). We found six structures showing NSP3 PL-Pro bound to ubiquitin domains from UBB, UBC, or UBA52 (a.k.a. ubiquitin-60S ribosomal protein L40). Four additional structures showed binding to the ubiquitin-like domains of ISG15 (a.k.a. interferon stimulated gene 15), which attaches to newly synthesized proteins; ISGylation does not induce degradation but likely disturbs virion assembly (22). Together, these 10 structures show in detail how NSP3 reverses either ubiquitination or ISGylation (Fig. 4B); a further two structures show how NSP3 may be blocked with inhibitors (Fig. 2).

Spike glycoprotein is known to bind receptors on the host cell membrane, thereby initiating membrane fusion and viral entry (33). We found 16 matching structures showing hijacking of ACE2, a carboxypeptidase that cleaves vasoactive peptides. One of these structures also shows binding to SLC6A19 (a.k.a. B^0^AT1), an amino acid transporter reported to act with ACE2 (57). Most structures show only short fragments of this protein; however, by consolidating several structures and sequence features, we created an integrated view showing the cellular context of the pre-fusion event, when spike first binds the ACE + SLC6A19 complex (Fig. 4B). Three further matching structures showed interactions between spike and host membrane receptor DPP4 (a.k.a. CD26), an aminopeptidase that SARS-CoV-2 may use to enter host immune cells (58). If so, DPP4 and ACE2 may interact with spike at distinct but overlapping binding sites (Fig. 4B) that may be useful as therapeutic targets. Some reports suggest that, although DPP4 may not be involved (59), SARS-CoV-2 directly infects some immune cells (60); if so, it may switch to a non-reproductive mode, leading to immune dysregulation (61) and apoptosis — as reported for SARS-CoV (62) and MERS-CoV (63) — thereby contributing to lymphopenia (64) and necrosis of lymphoid tissue (65) seen in severe COVID-19. We speculate that this cell-type-dependent state switching may be driven by NSP3 modifications of either ADP ribosylation, ubiquitination, or ISGlyation, and may involve the state switches noted above for NSP13 and nucleocapsid protein.

### Teams

We found structural evidence for interaction between only six SARS-CoV-2 proteins (Figs. 2 & 5); they divided into two distinct teams, each with three proteins.

Team 1 comprised NSP7, NSP8, and NSP12, all members of the viral RNA replication complex (Fig. 5A). NSP12 alone can replicate RNA, as can NSP7 + NSP8 acting together (25). However, replication is greatly stimulated by cooperative interactions between these proteins (66). Most combinations of these proteins were seen amongst the 60 matching structures found (Fig. 5A), although all models for NSP12 alone or with RNA derived from very remote viruses (e.g., enterovirus). The matrix of combinations (Fig. 5A) gives insight into how the complex may assemble, and shows that several proteins that may be involved in genome replication (29) are missing (NSP3, NSP9, NSP10, NSP13, NSP14, and NSP16). These outcomes demonstrate the value of modelling all available structural states, rather than only one or few states per protein.

Team 2 comprised NSP10, NSP14, and NSP16 (Fig. 5B). All matching structures found for NSP14 and NSP16 showed binding with NSP10, consistent with the belief that NSP10 is required for NSP16 RNA-cap methyltransferase activity (67), and also for NSP14 methyltransferase and exoribonuclease activities (68). In addition, NSP10 was found to form a homododecamer; however, these three oligomeric states of NSP10 share a common binding region (69) (Fig. 5B). This suggests NSP10, NSP14, and NSP16 interact competitively — in contrast to cooperative interactions seen in team 1. We speculate that NSP10 is produced at higher abundance than NSP14 or NSP16, or may be rate limiting for viral replication.

Finally, it is noteworthy that no interactions were found between the 12 virion or accessory proteins (Fig. 1, bottom third). This, again, highlights limitations in currently available structural data.

### Suspects

This leaves 18 of the 27 viral proteins in a final category we call suspects: these are proteins thought to play key roles in infection, but having no structural evidence of interaction with RNA, DNA, or other proteins (viral or human). We divided the suspects into two groups, based on matching structures.

Group 1 proteins were those with at least one matching structure: NSP1, NSP2, NSP4, NSP9, NSP15, E, ORF7a, and ORF9a. Some of these have been well studied (e.g., NSP5 had 256 matching structures). Yet none of these proteins had significant similarity to any experimentally determined 3D structure involving human proteins, or to any structure showing interactions between viral proteins — based on the methods used in this work.

Group 2 proteins were those with no matching structures: NSP6, ORF3a, matrix glycoprotein, ORF6, ORF8, and ORF9c, and ORF10. As noted previously, NSP2 could, arguably, be included in this group. These are structurally dark proteins (13), meaning not only is their structure unknown, but also that they have no significant sequence similarity to any experimentally determined 3D structure — based on the methods used in this work. These proteins are ripe candidates for advanced modelling strategies, e.g., using predicted residue contacts combined with deep learning (3).

### Conclusions

We have assembled a wealth of structural data, not available elsewhere, about the viral proteome. We used these data to construct a structural coverage map summarizing what is known — and not known — about the 3D structure of SARS-CoV-2 proteins. The coverage map also summarizes structural evidence for viral protein interaction teams and for mimicry and hijacking of host proteins — and identifies suspect viral proteins, with no evidence of interactions to other macromolecules. The map can be used by researchers to help find 3D models of interest, which can then be mapped with a large set of sequence features to provide insights into protein function. The resulting Aquaria-COVID resource (https://aquaria.ws/covid) aims to fulfil a vital role during the COVID-19 pandemic, helping scientists use emerging structural data to understand the molecular mechanisms underlying coronavirus infection.

Based on insights gained using our resource, we propose mechanisms by which SARS-CoV-2 may sense the type of cells it infects (via posttranslational modifications by NSP3) and, in certain cells, may switch focus from viral reproduction to disruption of host immunity (via hijacking by NSP13). Our resource lets researchers easily explore and assess the evidence for these mechanisms — or for others they may propose themselves — thereby helping direct future research.

## Materials and Methods

### SARS-CoV-2 Sequences

This study was based on the 14 protein sequences provided in UniProtKB/Swiss-Prot version 2020_03 (released April 22, 2020; https://www.uniprot.org/statistics/) as comprising the SARS-CoV-2 proteome. Swiss-Prot provides polyproteins 1a and 1ab (a.k.a. PP1a and PP1ab) as two separate entries, both identical for the first 4401 residues; PP1a then has four additional residues (‘GFAV’) not in PP1ab, which has 2695 additional residues not in PP1a. Swiss-Prot also indicates residue positions at which the polyproteins become cleaved into protein fragments, named NSP1 though NSP16. The NSP11 fragment comprises the last 13 residues of PP1a (4393–4405). The first 9 residues of NSP12 are identical to the first nine of NSP11, but the rest of that 919 residue long protein continues with a different sequence due to a functionally important frameshift between ORF1a and ORF1b (70). Thus, following cleavage, the proteome comprises a final total of 27 separate proteins.

### Sequence-to-Structure Alignments

The 14 SARS-CoV-2 sequences were then systematically compared with sequences derived from all known 3D structures from all organisms, based on PDB released on May 30, 2020. These comparisons used the latest version of HHblits (8) — an alignment method employing iterative comparisons of hidden Markov models (HMMs) — and using the processing pipeline defined previously (9), accepting all sequence-to-structure alignments with a significance threshold of E ≤ 10^−10^. The result is a database of protein sequence-to-structure homologies called PSSH2, which is used in the Aquaria web resource (9).

HHblits has been substantially updated since we last assessed the specificity and sensitivity of PSSH2 (9), therefore we repeated our assessment of resulting alignments, this time using CATH (16) for the determination of accuracy and precision. Our test data set comprised 23,028 sequences from the CATH nr40 data set. We built individual sequence profiles against UniClust30 and used these profiles to search against “PDB_full”, a database of HMMs for all PDB sequences. We then evaluated how many false positives were retrieved at an E-value lower than 10^−10^, where a false positive was seen to be a structure with a different CATH code at the level of homologous superfamily (H) or topology (T). We compared the ratio of false positives received with HH-suite3 and UniClust30 with a similar analysis for data produced in 2017 with HH-suite2 and UniProt20, and found that in both cases the false positive rate was at 2.5% at the homology level (H), and 1.9% at the topology level (T). The recovery rate, i.e. the ratio of proteins from the CATH nr40 data (with less than 40% sequence identity) found by our method that have the same CATH code, was slightly higher with HH-suite3 (8) (20.8% vs. 19.4%).

For each sequence-to-structure alignment derived from HHblits, the Aquaria interface shows the pairwise sequence identity score, thus providing an intuitive indication of how closely related the given region of SARS-CoV-2 is to the sequence of the matched structure. However, to more accurately assess the quality of the match, Aquaria also gives an E-value, calculated by comparing two HMMs, one generated for each of these two sequences.

### PredictProtein Features

To facilitate analysis of SARS-CoV-2 sequences, we enhanced the Aquaria resource to include PredictProtein features (18), thus providing a very rich set of predicted features for all Swiss-Protein sequences. The four PredictProtein feature sets used in this work were fetched via:

https://api.predictprotein.org/v1/results/molart/:uniprot_id

#### Conservation

The first PredictProtein feature set is generated by ConSurf (71) and gives, for each residue, a score between 1 and 9, corresponding to very low and very high conservation, respectively. These scores estimate the evolutionary rate in protein families, based on evolutionary relatedness between the query protein and its homologues from UniProt using empirical Bayesian methods (72).

#### Disorder

This feature set gives consensus predictions generated by Meta-Disorder (73), which combines outputs of several structure-based disorder predictors to classify each residue as either disordered or not disordered.

#### Flexibility

This feature set predicts, for each residue, normalized B-factor values expected to be observed in an X-ray-derived structure, generated by PROFbval (73). For each residue, PROFbval provides a score between 0 and 100; a score of 50 indicates average flexibility, while ≥71 indicates highly flexible residues.

#### Topology

This feature set is generated by TMSEG (74), a machine learning model that uses evolutionary-derived information to predict regions of a protein that traverse membranes, as well as the subcellular locations of complementary (non-transmembrane) regions.

### SNAP2 Features

We further enhanced Aquaria to include SNAP2 (17) features, which provide information on the mutational propensities for each residue position. In Aquaria, two SNAP2 features sets are fetched via:

https://rostlab.org/services/aquaria/snap4aquaria/json.php?uniprotAcc=:uniprot_id

#### Mutational Sensitivity

The first SNAP2 feature set provides, for each residue position, a list of 20 scores that indicate the predicted functional consequences of the position being occupied by each of the 20 standard amino acids. Large, positive scores (up to 100) indicate substitutions likely to have deleterious changes, while negative scores (down to -100) indicate no likely functional change. From these 20 values, a single summary score is calculated based on the total fraction of substitutions predicted to have deleterious effect, taken to be those with a score > 40. The summary scores are used to generate a red to blue color map, indicating residues with highest to least functional importance, respectively.

#### Mutational Score

The second SNAP2 feature set is based on the same 20 scores above, but calculates the single summary score for each residue as the average of the individual scores for each of the 20 standard amino acids.

### UniProt Features

UniProt features are curated annotations, and therefore largely complement the automatically generated PredictProtein features. In Aquaria, for each protein sequence, the UniProt feature collection is fetched via:

https://www.uniprot.org/uniprot/:uniprot_id.xml

### CATH Features

For this work, we further enhanced Aquaria to include CATH domain annotations (16). For most protein sequences, Aquaria fetches two CATH feature sets via APIs given at:

https://github.com/UCLOrengoGroup/cath-api-docs

For SARS-CoV-2 proteins, however, CATH annotations are not yet fully available via the above APIs. In this work, we used a pre-release version of these annotations, derived by scanning the UniProt pre-release sequences against the CATH-Gene3D v4.3 FunFams HMM library (75).

Domain assignments were obtained using cath-resolve-hits and curated manually (75). For the SARS-CoV-2 sequences, Aquaria fetches two CATH feature sets via:

https://aquaria.ws/covid19cath/P0DTC2

#### Superfamilies

The first CATH feature set identifies regions of protein sequences across a wide variety of organisms that are expected to have very similar 3D structures and to have general biological functions in common.

#### Functional Families

Also known as FunFams, this feature set partitions each superfamily into subsets expected to have more specific biological functions in common (75).

When examining a specific superfamily or functional family domain in Aquaria, the browser uses additional CATH API endpoints (via the link above) to create compact, interactive data visualizations that give access to detailed information on the biological function and phylogenetic distribution of proteins containing that domain.

### Average Feature Scores

In the results, we include average scores for some of the above feature sets. The average scores given for NSP3 regions were calculated using features provided for the PP1a sequence; however, the average whole-protein scores given for each of the 15 PP1ab proteins were calculated using features provided for the PP1ab sequence.

### Aquaria Core

For this work, the Aquaria core codebase has been substantially refactored. We changed all client-server data exchanges to use get requests, instead of sockets as used previously (9). This allowed us to further implement both server- and client-side caching, resulting in greatly improved performance. Another major change was to the user interface, where we removed the previously used Java applet (9), and replaced it with Jolecule, a JavaScript-based molecular graphics component (https://jolecule.com/), that we further augmented to enable feature mapping. In this work, Jolecule was used to determine the set of residues comprising intermolecular contacts by selecting all residues for one protein, then applying Jolecule’s ‘Neighbours’ function. This highlights all residues in which any atom is within 5 Å.

### Structure Coverage Matrix

We created a web page featuring a matrix view of the 14 UniProt sequence, and showing the total number of matching structures found. This page allows navigation to the corresponding Aquaria page for each protein sequence, from where all matching structures can be seen and mapped with sequence features.

### Structural Coverage Map

We created an additional web page using a layout derived from the organization of the viral genome. For each contiguous region of the viral proteome that had matching structures, we selected a single representative structure (Fig. 2), primarily based on identity to the SARS-CoV-2 sequence. However, in some cases, representatives were chosen that best illustrated the consensus biological assembly seen across all matching structures, or showed the simplest assembly. Under the name of each viral protein, the total number of matching structures found in PSSH2 (9) is indicated. Below each structure, a tree graph is drawn where there is structural evidence of mimicry (i.e., where the viral sequence aligns onto human proteins), hijacking (i.e., where the matching structures show binding between viral and human proteins, or with DNA, RNA, antibodies, or inhibitory factors), or physical interaction with other viral proteins. When these tree graphs are missing, none of the matching structures meet these criteria.

## Acknowledgements

Thanks to Tim Mercer and Giulia Wang (Garvan Institute, Australia), Phil Austin (University of Sydney, Australia) and Lucy van Dorp (UCL Genetics Institute, UK) for helpful feedback and discussions, to Ian Sillitoe (UCL, UK) for helpful advice regarding the CATH API, to Tim Karl, Michael Bernhofer, and Maria Littmann (TUM, Germany) for advice regarding the PredictProtein server. We are grateful to Max Ott (CSIRO, Australia) for advice on improving the performance and reliability of the Aquaria web application.

## Abbreviations

ADPr: (ADP-ribose)
BtCoV-HKU4: (bat coronavirus HKU4)
CHIKV: (Chikungunya virus)
FCoV: (feline coronavirus)
FMDV: (foot-and-mouth disease virus)
HMM: (hidden Markov model)
IBV: (avian infectious bronchitis virus)
ISG15: (interferon stimulated gene 15)
MHV-A59: (mouse hepatitis virus A59)
PDB: (Protein Data Bank) PP1a (polyprotein 1a)
PP1ab: (polyprotein 1ab)
SARS-CoV: (Severe Acute Respiratory Syndrome Coronavirus)
Ubl: (ubiquitin-like)

## Author Contributions

SIOD designed this study and led in coordinating co-author contributions, in data analysis, in manuscript writing, and in figure preparation. He also participated in the database generation. AS led in 3D model generation, in PSSH2 validation, and was involved in data analysis and in writing the manuscript. NS coordinated integration of Aquaria code improvements and was involved in data analysis, in writing the manuscript, and in creating the tables. CS led in the design of graphical user interface elements and contributed to figure generation. SK led in implementing CATH, SNAP2, and PredictProtein features into Aquaria, participated in data analysis, and assisted in creating the tables. BKH led the integration for Jolecule into the Aquaria user interface. SA and MA implemented key new features to the molecular graphics components. JP contributed to data analysis and provided strategic input into the manuscript. CD helped with integration of SNAP2 and PredictProtein features into Aquaria, and was involved in data analysis, in writing the manuscript, and in creating the tables. NB participated with the implementation of CATH features into Aquaria and assisted in creating the tables. BR contributed to model generation, oversaw SNAP2 and PredictProtein developments, and gave strategic input into the manuscript.

## Ethics Declarations

The authors declare no competing interests.

## Supplementary Information

[SupplementaryInformation](http://docs.google.com/document/d/1DSOtarLraNL8yGvk54-GkNIVw34wa2ssMS8OYhKaIV0/edit#)

## Data Availability

All data and code will be made publicly available.

## Supplemental Information

**Table S1:** SARS-CoV-2 protein sequences used in this study

| Sequence | Source | Identifier |
| --- | --- | --- |
| Polyprotein 1a | UniProt | P0DTC1 |
| Polyprotein 1ab | UniProt | P0DTD1 |
| Spike glycoprotein | UniProt | P0DTC2 |
| ORF3a protein | UniProt | P0DTC3 |
| Envelope protein | UniProt | P0DTC4 |
| Matrix glycoprotein | UniProt | P0DTC5 |
| ORF6 protein | UniProt | P0DTC6 |
| ORF7a protein | UniProt | P0DTC7 |
| ORF7b protein | UniProt | P0DTD8 |
| ORF8 protein | UniProt | P0DTC8 |
| Nucleocapsid protein | UniProt | P0DTC9 |
| ORF9b protein | UniProt | P0DTD2 |
| ORF9c protein | UniProt | P0DTD3 |
| ORF10 protein | UniProt | A0A663DJA2 |

**Table S2:**
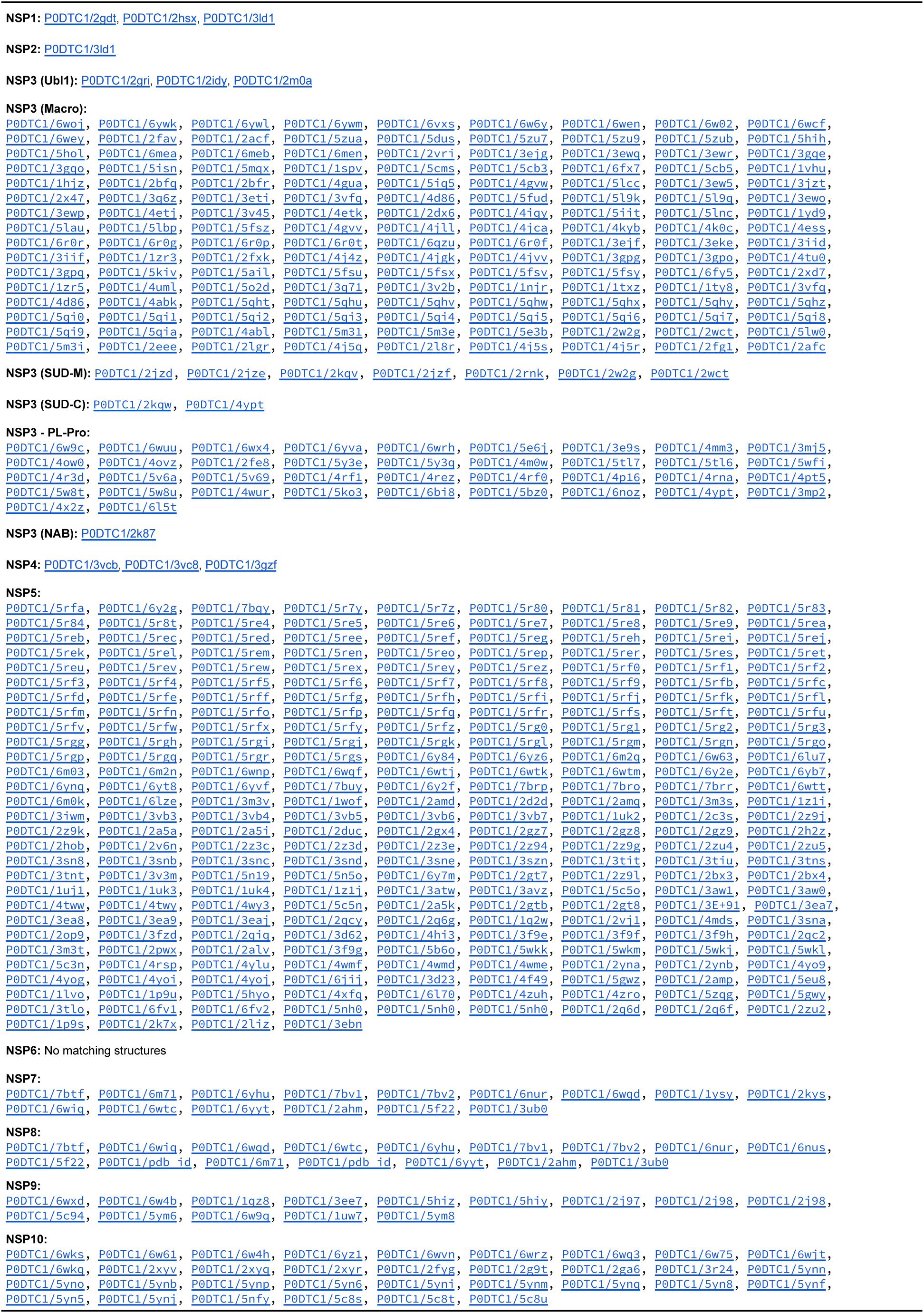
Matching structures for SARS-CoV-2 polyprotein 1a

**Table S3:**
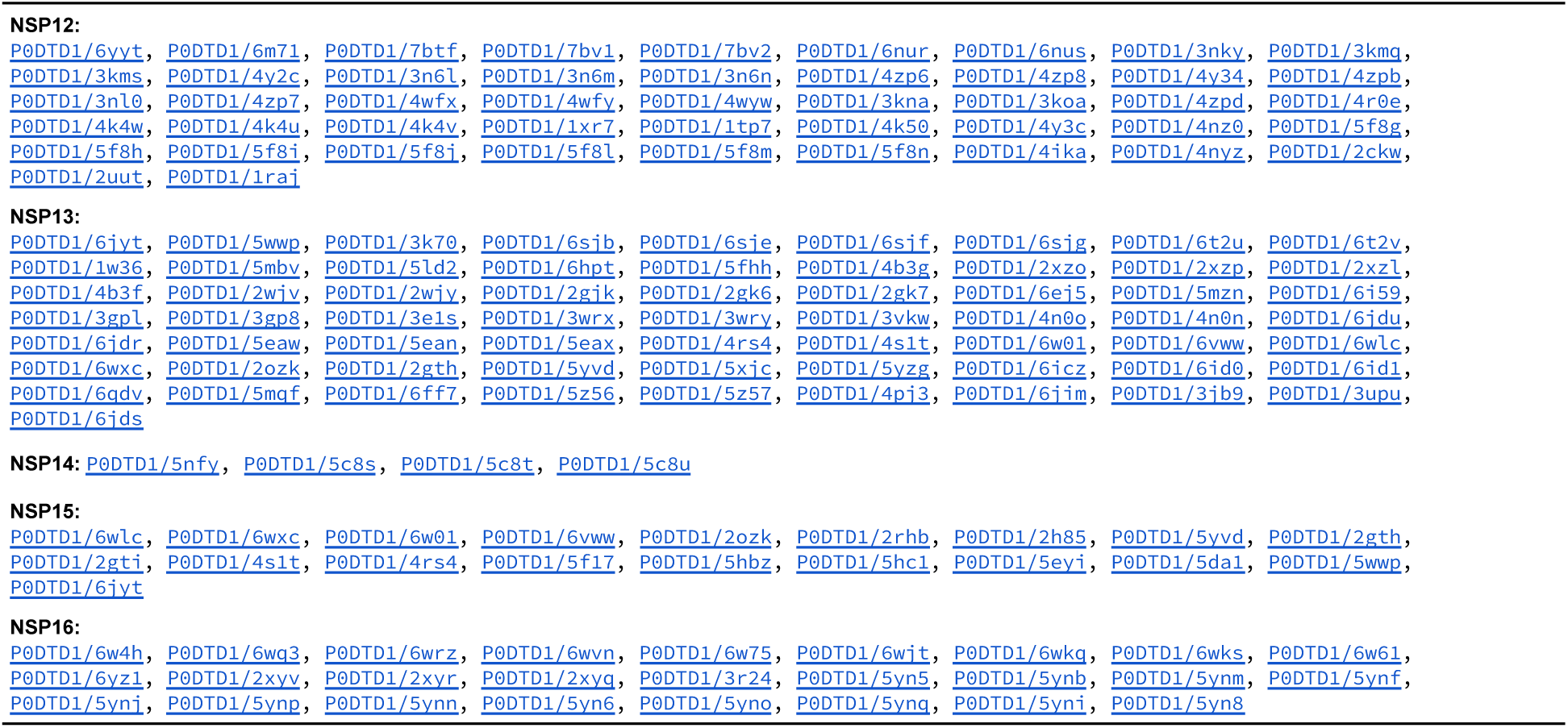
Matching structures for SARS-CoV-2 polyprotein 1b

**Table S4:**
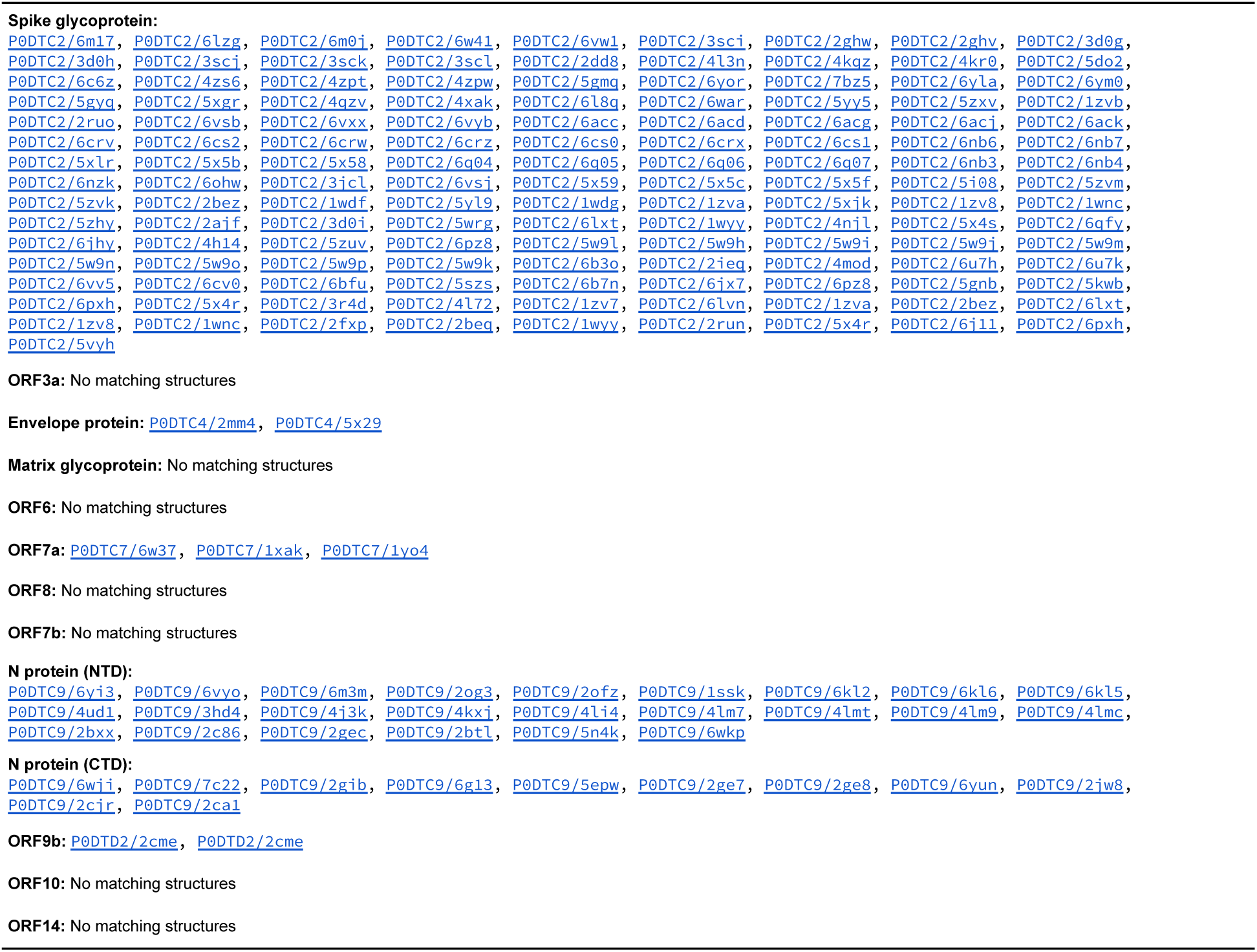
Matching structures for SARS-CoV-2 Virion and Accessory Proteins

**Table S5:**
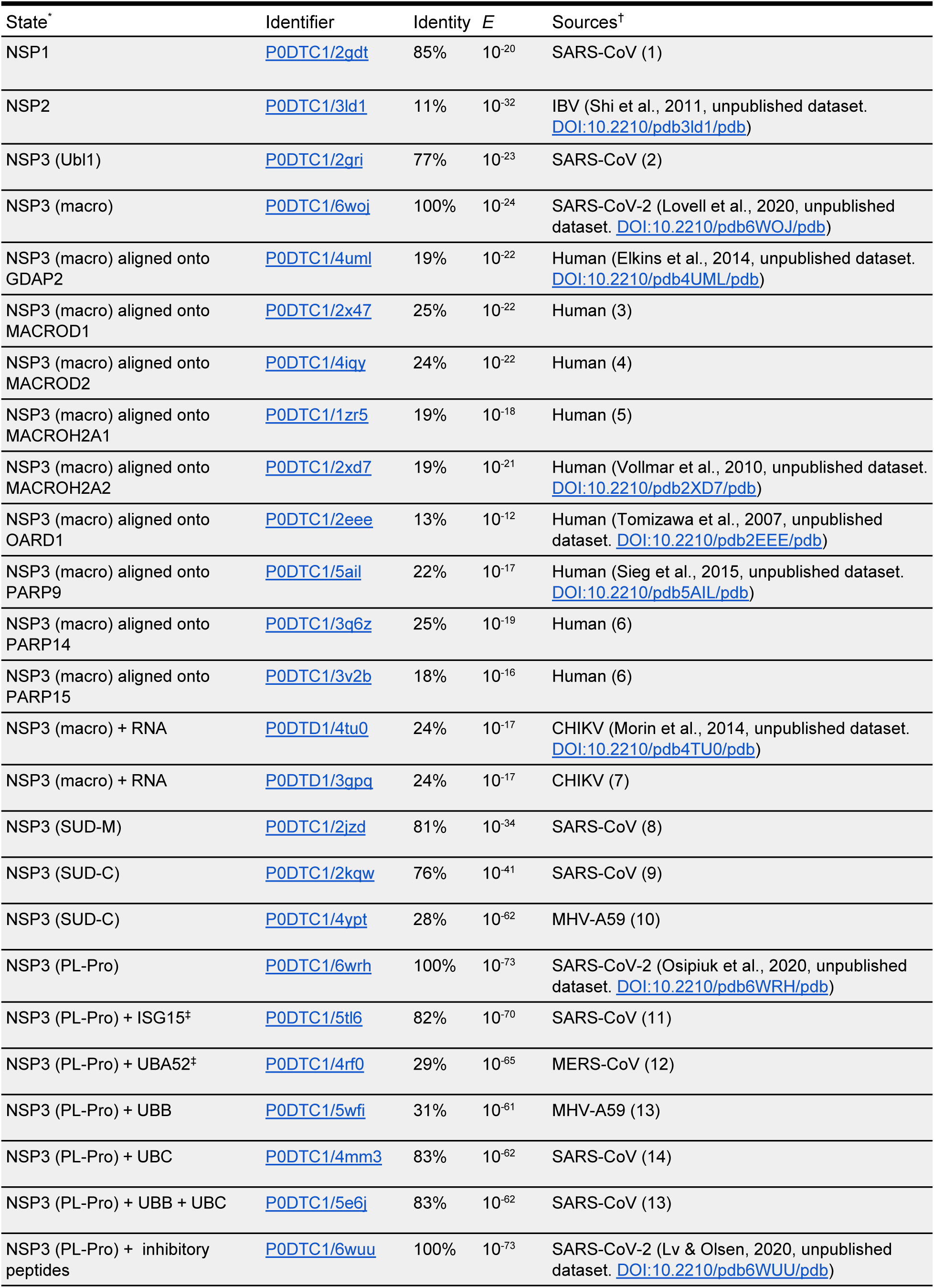
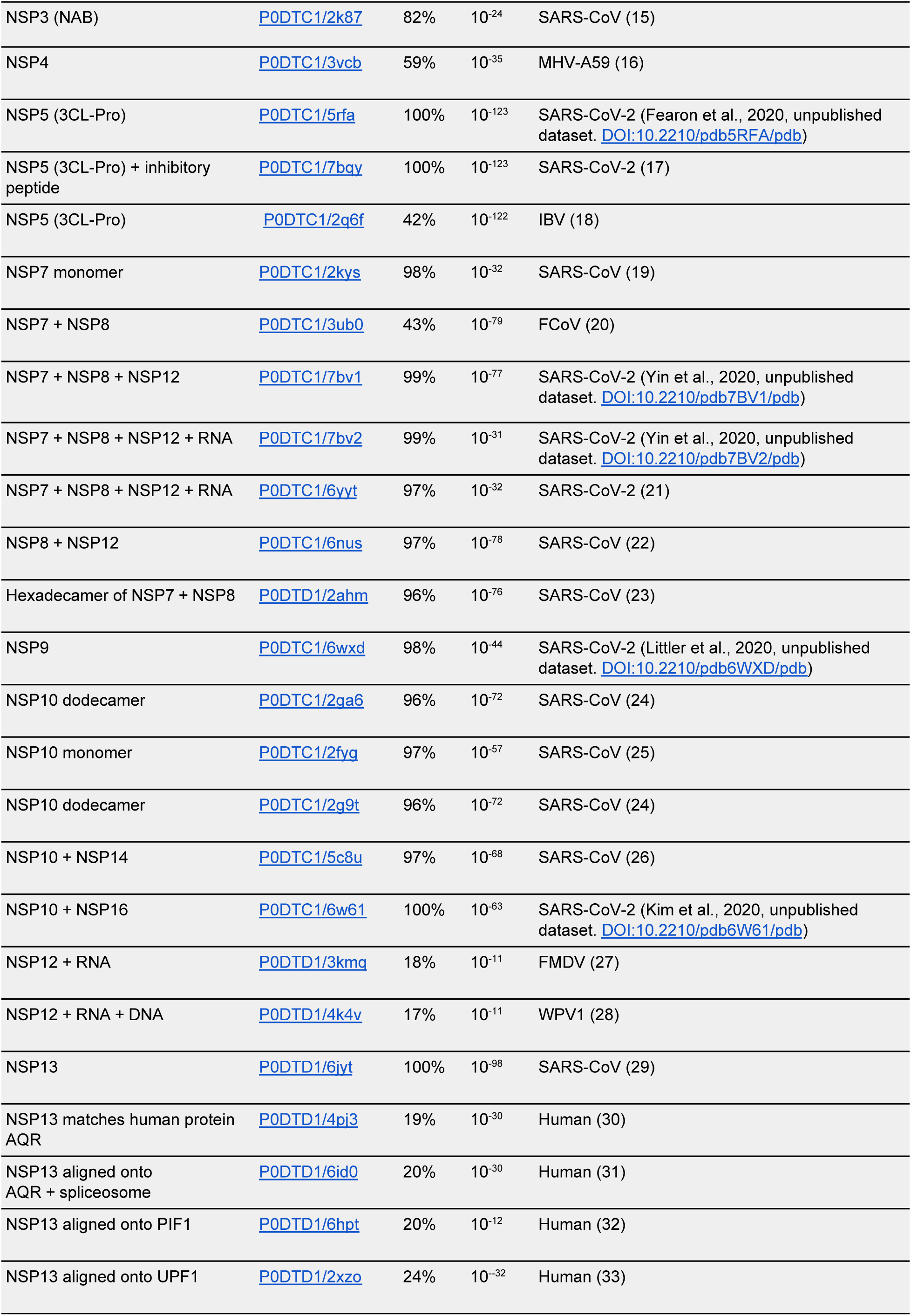
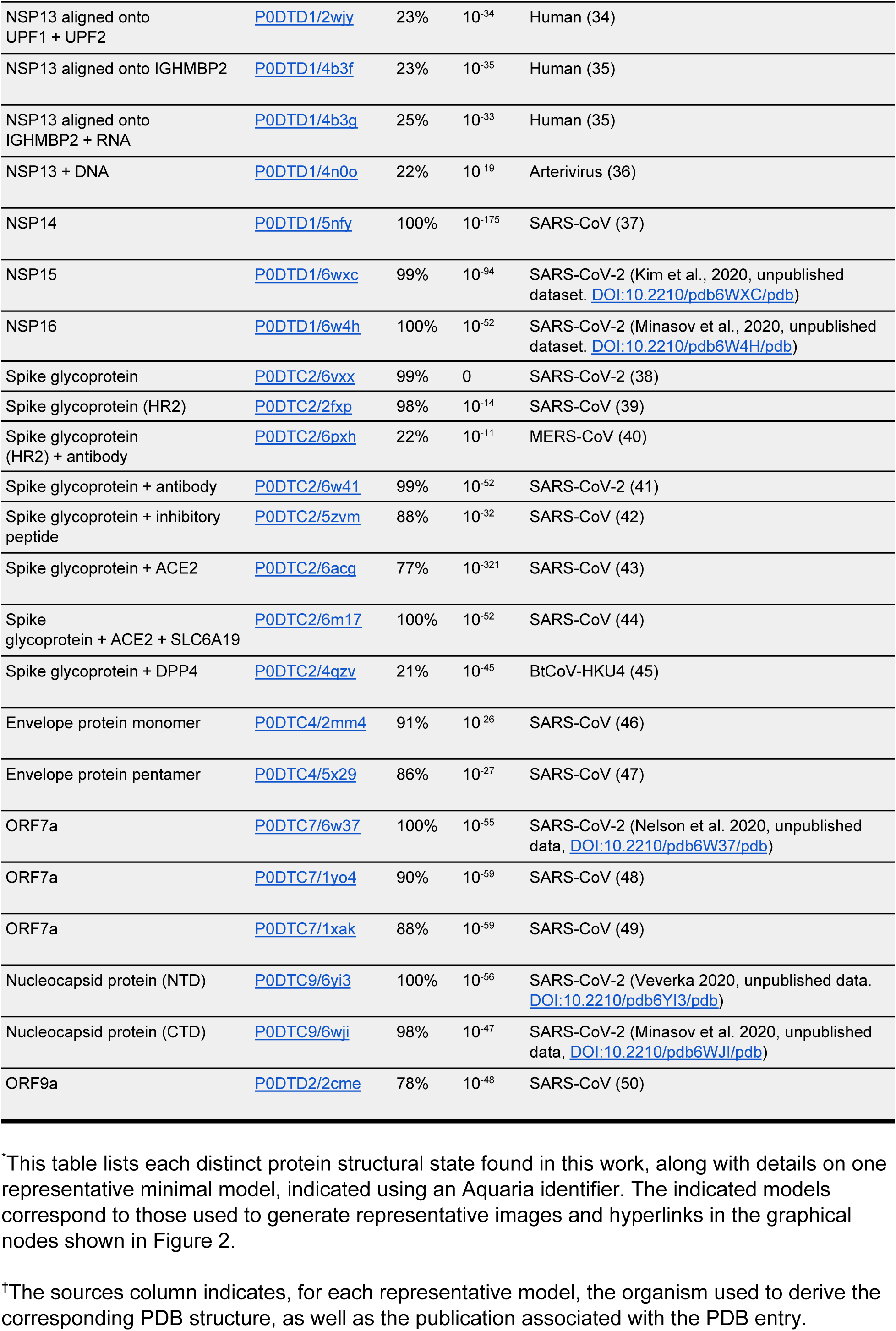
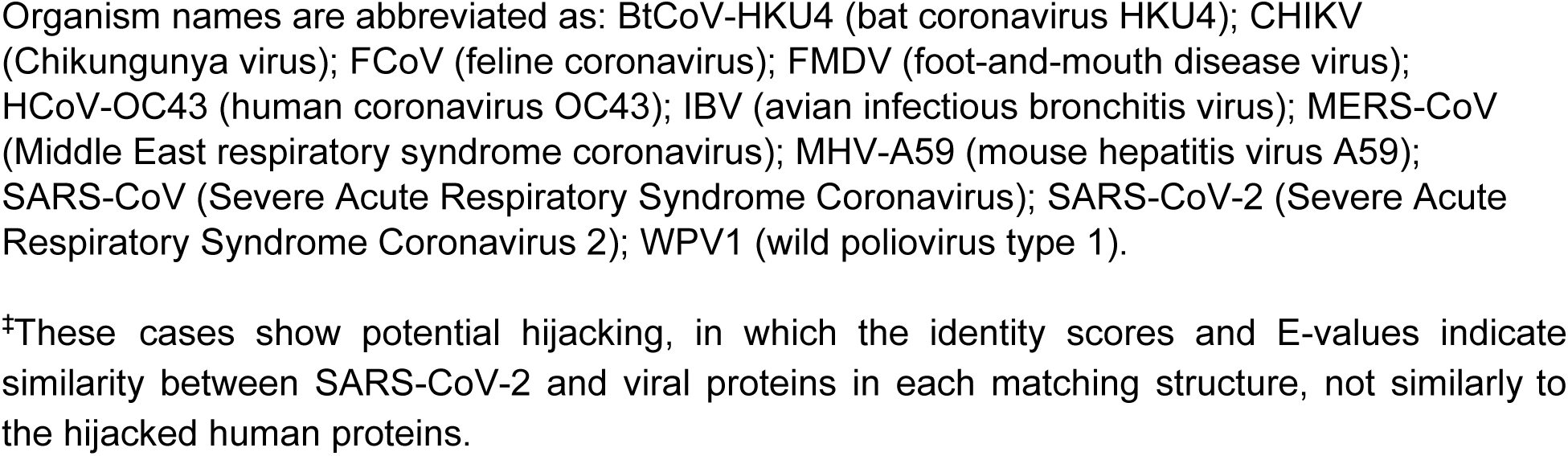
SARS-CoV-2 minimal models used in Figure 2

**Table S6:**
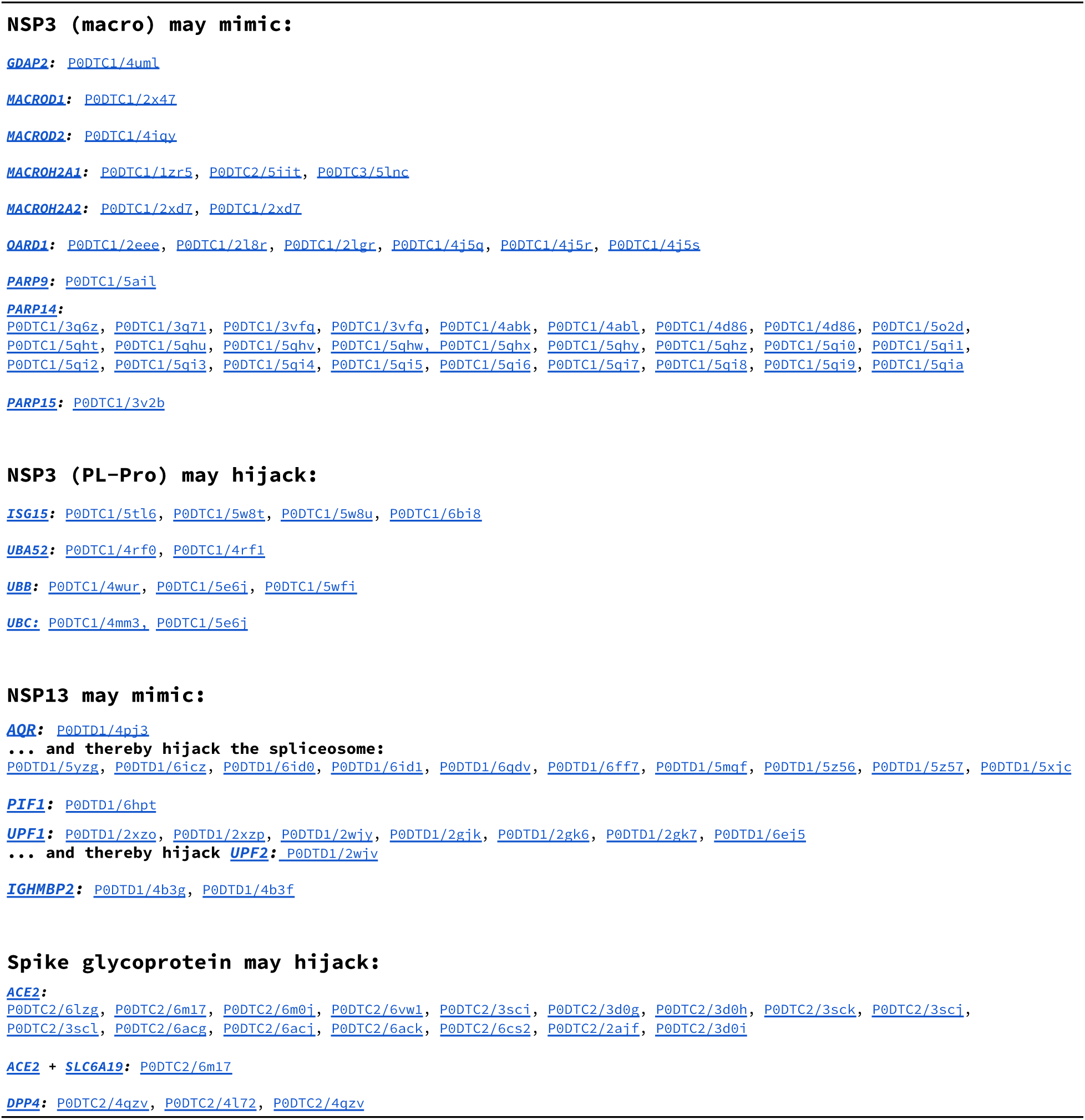
Structural evidence for mimicry and hijacking of human proteins

## References

1. Berman HM, Westbrook J, Feng Z, Gilliland G, Bhat TN, Weissig H, et al. The Protein Data Bank. Nucleic Acids Res. 2000 Jan 1;28(1):235–42.

2. Jaimes JA, André NM, Chappie JS, Millet JK, Whittaker GR. Phylogenetic Analysis and Structural Modeling of SARS-CoV-2 Spike Protein Reveals an Evolutionary Distinct and Proteolytically Sensitive Activation Loop. J Mol Biol. 2020 May 1;432(10):3309–25.

3. Senior AW, Evans R, Jumper J, Kirkpatrick J, Sifre L, Green T, et al. Improved protein structure prediction using potentials from deep learning. Nature. 2020 Jan;577(7792):706–10.

4. Zheng W, Li Y, Zhang C, Pearce R, Mortuza SM, Zhang Y. Deep-learning contact-map guided protein structure prediction in CASP13. Proteins. 2019;87(12):1149–64.

5. Rohl CA, Strauss CEM, Misura KMS, Baker D. Protein Structure Prediction Using Rosetta. In: Methods in Enzymology [Internet]. Academic Press; 2004 [cited 2020 Jul 21]. p. 66–93. (Numerical Computer Methods, Part D; vol. 383). Available from: http://www.sciencedirect.com/science/article/pii/S0076687904830040

6. Waterhouse A, Bertoni M, Bienert S, Studer G, Tauriello G, Gumienny R, et al. SWISS-MODEL: homology modelling of protein structures and complexes. Nucleic Acids Res. 2018 Jul 2;46(W1):W296–303.

7. Heo L, Feig M. Modeling of Severe Acute Respiratory Syndrome Coronavirus 2 (SARS-CoV-2) Proteins by Machine Learning and Physics-Based Refinement. bioRxiv. 2020 Mar 28;2020.03.25.008904.

8. Steinegger M, Meier M, Mirdita M, Vöhringer H, Haunsberger SJ, Söding J. HH-suite3 for fast remote homology detection and deep protein annotation. BMC Bioinformatics. 2019 Sep 14;20(1):473.

9. O’Donoghue SI, Sabir KS, Kalemanov M, Stolte C, Wellmann B, Ho V, et al. Aquaria: simplifying discovery and insight from protein structures. Nat Methods. 2015 Feb;12(2):98–9.

10. Heinrich J, Kaur S, O’Donoghue S. Evaluating the Effectiveness of Color to Convey Alignment Quality in Macromolecular Structures. In Hobart, Australia: IEEE; 2015 [cited 2020 Jul 21]. Available from: https://ieeexplore.ieee.org/document/7314292

11. O’Donoghue SI, Baldi BF, Clark SJ, Darling AE, Hogan JM, Kaur S, et al. Visualization of Biomedical Data. Annu Rev Biomed Data Sci. 2018 Jul 20;1(1):275–304.

12. The UniProt Consortium. UniProt: a worldwide hub of protein knowledge. Nucleic Acids Res. 2019 Jan 8;47(D1):D506–15.

13. Perdigão N, Heinrich J, Stolte C, Sabir KS, Buckley MJ, Tabor B, et al. Unexpected features of the dark proteome. Proc Natl Acad Sci. 2015 Dec 29;112(52):15898–903.

14. Elde NC, Malik HS. The evolutionary conundrum of pathogen mimicry. Nat Rev Microbiol. 2009 Nov;7(11):787–97.

15. Davey NE, Travé G, Gibson TJ. How viruses hijack cell regulation. Trends Biochem Sci. 2011 Mar;36(3):159–69.

16. Dawson NL, Lewis TE, Das S, Lees JG, Lee D, Ashford P, et al. CATH: an expanded resource to predict protein function through structure and sequence. Nucleic Acids Res. 2017 Jan 4;45(D1):D289–95.

17. Hecht M, Bromberg Y, Rost B. Better prediction of functional effects for sequence variants. BMC Genomics. 2015 Jun 18;16(8):S1.

18. Yachdav G, Kloppmann E, Kajan L, Hecht M, Goldberg T, Hamp T, et al. PredictProtein—an open resource for online prediction of protein structural and functional features. Nucleic Acids Res. 2014 Jul 1;42(W1):W337–43.

19. Kamitani W, Huang C, Narayanan K, Lokugamage KG, Makino S. A two-pronged strategy to suppress host protein synthesis by SARS coronavirus Nsp1 protein. Nat Struct Mol Biol. 2009 Nov;16(11):1134–40.

20. Almeida MS, Johnson MA, Herrmann T, Geralt M, Wüthrich K. Novel β-Barrel Fold in the Nuclear Magnetic Resonance Structure of the Replicase Nonstructural Protein 1 from the Severe Acute Respiratory Syndrome Coronavirus. J Virol. 2007 Apr 1;81(7):3151–61.

21. Cornillez-Ty CT, Liao L, Yates JR, Kuhn P, Buchmeier MJ. Severe Acute Respiratory Syndrome Coronavirus Nonstructural Protein 2 Interacts with a Host Protein Complex Involved in Mitochondrial Biogenesis and Intracellular Signaling. J Virol. 2009 Oct 1;83(19):10314–8.

22. Lei J, Kusov Y, Hilgenfeld R. Nsp3 of coronaviruses: Structures and functions of a large multi-domain protein. Antiviral Res. 2018 Jan;149:58–74.

23. Angelini MM, Neuman BW, Buchmeier MJ. Untangling Membrane Rearrangement in the Nidovirales. DNA Cell Biol. 2014 Mar;33(3):122–7.

24. Xu X, Lou Z, Ma Y, Chen X, Yang Z, Tong X, et al. Crystal Structure of the C-Terminal Cytoplasmic Domain of Non-Structural Protein 4 from Mouse Hepatitis Virus A59. Masucci MG, editor. PLoS ONE. 2009 Jul 10;4(7):e6217.

25. te Velthuis AJW, van den Worm SHE, Snijder EJ. The SARS-coronavirus nsp7+nsp8 complex is a unique multimeric RNA polymerase capable of both de novo initiation and primer extension. Nucleic Acids Res. 2012 Feb;40(4):1737–47.

26. Miknis ZJ, Donaldson EF, Umland TC, Rimmer RA, Baric RS, Schultz LW. Severe Acute Respiratory Syndrome Coronavirus nsp9 Dimerization Is Essential for Efficient Viral Growth. J Virol. 2009 Apr 1;83(7):3007–18.

27. Bouvet M, Imbert I, Subissi L, Gluais L, Canard B, Decroly E. RNA 3’-end mismatch excision by the severe acute respiratory syndrome coronavirus nonstructural protein nsp10/nsp14 exoribonuclease complex. Proc Natl Acad Sci. 2012 Jun 12;109(24):9372–7.

28. Yin W, Mao C, Luan X, Shen D-D, Shen Q, Su H, et al. Structural basis for inhibition of the RNA-dependent RNA polymerase from SARS-CoV-2 by remdesivir. Science. 2020 Jun 26;368(6498):1499–504.

29. Subissi L, Imbert I, Ferron F, Collet A, Coutard B, Decroly E, et al. SARS-CoV ORF1b-encoded nonstructural proteins 12–16: Replicative enzymes as antiviral targets. Antiviral Res. 2014 Jan;101:122–30.

30. Minskaia E, Hertzig T, Gorbalenya AE, Campanacci V, Cambillau C, Canard B, et al. Discovery of an RNA virus 3’->5’ exoribonuclease that is critically involved in coronavirus RNA synthesis. Proc Natl Acad Sci. 2006 Mar 28;103(13):5108–13.

31. Ricagno S, Egloff M-P, Ulferts R, Coutard B, Nurizzo D, Campanacci V, et al. Crystal structure and mechanistic determinants of SARS coronavirus nonstructural protein 15 define an endoribonuclease family. Proc Natl Acad Sci. 2006 Aug 8;103(32):11892–7.

32. Bouvet M, Debarnot C, Imbert I, Selisko B, Snijder EJ, Canard B, et al. In Vitro Reconstitution of SARS-Coronavirus mRNA Cap Methylation. Buchmeier MJ, editor. PLoS Pathog. 2010 Apr 22;6(4):e1000863.

33. Hoffmann M, Kleine-Weber H, Schroeder S, Krüger N, Herrler T, Erichsen S, et al. SARS-CoV-2 Cell Entry Depends on ACE2 and TMPRSS2 and Is Blocked by a Clinically Proven Protease Inhibitor. Cell. 2020 Apr;181(2):271-280.e8.

34. Lu W, Zheng B-J, Xu K, Schwarz W, Du L, Wong CKL, et al. Severe acute respiratory syndrome-associated coronavirus 3a protein forms an ion channel and modulates virus release. Proc Natl Acad Sci. 2006 Aug 15;103(33):12540–5.

35. Surya W, Li Y, Torres J. Structural model of the SARS coronavirus E channel in LMPG micelles. Biochim Biophys Acta BBA - Biomembr. 2018 Jun;1860(6):1309–17.

36. Vennema H, Godeke GJ, Rossen JW, Voorhout WF, Horzinek MC, Opstelten DJ, et al. Nucleocapsid-independent assembly of coronavirus-like particles by co-expression of viral envelope protein genes. EMBO J. 1996 Apr;15(8):2020–8.

37. Frieman M, Yount B, Heise M, Kopecky-Bromberg SA, Palese P, Baric RS. Severe Acute Respiratory Syndrome Coronavirus ORF6 Antagonizes STAT1 Function by Sequestering Nuclear Import Factors on the Rough Endoplasmic Reticulum/Golgi Membrane. J Virol. 2007 Sep 15;81(18):9812–24.

38. Taylor JK, Coleman CM, Postel S, Sisk JM, Bernbaum JG, Venkataraman T, et al. Severe Acute Respiratory Syndrome Coronavirus ORF7a Inhibits Bone Marrow Stromal Antigen 2 Virion Tethering through a Novel Mechanism of Glycosylation Interference. García-Sastre A, editor. J Virol. 2015 Dec 1;89(23):11820–33.

39. Schaecher SR, Mackenzie JM, Pekosz A. The ORF7b Protein of Severe Acute Respiratory Syndrome Coronavirus (SARS-CoV) Is Expressed in Virus-Infected Cells and Incorporated into SARS-CoV Particles. J Virol. 2007 Jan 15;81(2):718–31.

40. Li J-Y, Liao C-H, Wang Q, Tan Y-J, Luo R, Qiu Y, et al. The ORF6, ORF8 and nucleocapsid proteins of SARS-CoV-2 inhibit type I interferon signaling pathway. Virus Res. 2020 Sep;286:198074.

41. Grunewald ME, Fehr AR, Athmer J, Perlman S. The coronavirus nucleocapsid protein is ADP-ribosylated. Virology. 2018 Apr;517:62–8.

42. Surjit M, Kumar R, Mishra RN, Reddy MK, Chow VTK, Lal SK. The Severe Acute Respiratory Syndrome Coronavirus Nucleocapsid Protein Is Phosphorylated and Localizes in the Cytoplasm by 14-3-3-Mediated Translocation. J Virol. 2005 Sep 1;79(17):11476–86.

43. Shi C-S, Qi H-Y, Boularan C, Huang N-N, Abu-Asab M, Shelhamer JH, et al. SARS-Coronavirus Open Reading Frame-9b Suppresses Innate Immunity by Targeting Mitochondria and the MAVS/TRAF3/TRAF6 Signalosome. J Immunol. 2014 Sep 15;193(6):3080–9.

44. Meier C, Aricescu AR, Assenberg R, Aplin RT, Gilbert RJC, Grimes JM, et al. The Crystal Structure of ORF-9b, a Lipid Binding Protein from the SARS Coronavirus. Structure. 2006 Jul;14(7):1157–65.

45. Gordon DE, Jang GM, Bouhaddou M, Xu J, Obernier K, White KM, et al. A SARS-CoV-2 protein interaction map reveals targets for drug repurposing. Nature. 2020 Jul;583(7816):459–68.

46. Pan J, Peng X, Gao Y, Li Z, Lu X, Chen Y, et al. Genome-Wide Analysis of Protein-Protein Interactions and Involvement of Viral Proteins in SARS-CoV Replication. PLOS ONE. 2008 Oct 1;3(10):e3299.

47. O’Sullivan J, Tedim Ferreira M, Gagné J-P, Sharma AK, Hendzel MJ, Masson J-Y, et al. Emerging roles of eraser enzymes in the dynamic control of protein ADP-ribosylation. Nat Commun. 2019 Dec;10(1):1182.

48. Schäfer A, Baric R. Epigenetic Landscape during Coronavirus Infection. Pathogens. 2017 Feb 15;6(1):8.

49. Iwata H, Goettsch C, Sharma A, Ricchiuto P, Goh WWB, Halu A, et al. PARP9 and PARP14 cross-regulate macrophage activation via STAT1 ADP-ribosylation. Nat Commun. 2016 Nov;7(1):12849.

50. Varga Z, Flammer AJ, Steiger P, Haberecker M, Andermatt R, Zinkernagel AS, et al. Endothelial cell infection and endotheliitis in COVID-19. The Lancet. 2020 May;395(10234):1417–8.

51. Graham RL, Baric RS. Recombination, Reservoirs, and the Modular Spike: Mechanisms of Coronavirus Cross-Species Transmission. J Virol. 2010 Apr 1;84(7):3134–46.

52. Umate P, Tuteja N, Tuteja R. Genome-wide comprehensive analysis of human helicases. Commun Integr Biol. 2011;4(1):118–37.

53. Yu H-H, Chu K-H, Yang Y-H, Lee J-H, Wang L-C, Lin Y-T, et al. Genetics and Immunopathogenesis of IgA Nephropathy. Clin Rev Allergy Immunol. 2011 Oct;41(2):198–213.

54. Bauer G. The variability of the serological response to SARS-corona virus-2: Potential resolution of ambiguity through determination of avidity (functional affinity). J Med Virol. 2020 Jul 15;jmv.26262.

55. Wada M, Lokugamage KG, Nakagawa K, Narayanan K, Makino S. Interplay between coronavirus, a cytoplasmic RNA virus, and nonsense-mediated mRNA decay pathway. Proc Natl Acad Sci. 2018 Oct 23;115(43):E10157–66.

56. Aviv A. Telomeres and COVID-19. FASEB J. 2020 Jun;34(6):7247–52.

57. Camargo SMR, Singer D, Makrides V, Huggel K, Pos KM, Wagner CA, et al. Tissue-Specific Amino Acid Transporter Partners ACE2 and Collectrin Differentially Interact With Hartnup Mutations. Gastroenterology. 2009 Mar;136(3):872-882.e3.

58. Radzikowska U, Ding M, Tan G, Zhakparov D, Peng Y, Wawrzyniak P, et al. Distribution of ACE2, CD147, CD26, and other SARS-CoV-2 associated molecules in tissues and immune cells in health and in asthma, COPD, obesity, hypertension, and COVID-19 risk factors. Allergy. 2020 Aug 24;all.14429.

59. Tai W, He L, Zhang X, Pu J, Voronin D, Jiang S, et al. Characterization of the receptor-binding domain (RBD) of 2019 novel coronavirus: implication for development of RBD protein as a viral attachment inhibitor and vaccine. Cell Mol Immunol. 2020 Jun;17(6):613–20.

60. Li H, Liu L, Zhang D, Xu J, Dai H, Tang N, et al. SARS-CoV-2 and viral sepsis: observations and hypotheses. The Lancet. 2020 May;395(10235):1517–20.

61. Qin C, Zhou L, Hu Z, Zhang S, Yang S, Tao Y, et al. Dysregulation of Immune Response in Patients With Coronavirus 2019 (COVID-19) in Wuhan, China. Clin Infect Dis. 2020 Jul 28;71(15):762–8.

62. Wang H, Mao Y, Ju L, Zhang J, Liu Z, Zhou X, et al. Detection and Monitoring of SARS Coronavirus in the Plasma and Peripheral Blood Lymphocytes of Patients with Severe Acute Respiratory Syndrome. Clin Chem. 2004 Jul 1;50(7):1237–40.

63. Chu H, Zhou J, Wong BH-Y, Li C, Chan JF-W, Cheng Z-S, et al. Middle East Respiratory Syndrome Coronavirus Efficiently Infects Human Primary T Lymphocytes and Activates the Extrinsic and Intrinsic Apoptosis Pathways. J Infect Dis. 2016 Mar 15;213(6):904–14.

64. Tan L, Wang Q, Zhang D, Ding J, Huang Q, Tang Y-Q, et al. Lymphopenia predicts disease severity of COVID-19: a descriptive and predictive study. Signal Transduct Target Ther. 2020 Dec;5(1):33.

65. chen yongwen, Feng Z, Diao B, Wang R, Wang G, Wang C, et al. The Novel Severe Acute Respiratory Syndrome Coronavirus 2 (SARS-CoV-2) Directly Decimates Human Spleens and Lymph Nodes [Internet]. Infectious Diseases (except HIV/AIDS); 2020 Mar [cited 2020 Sep 23]. Available from: http://medrxiv.org/lookup/doi/10.1101/2020.03.27.20045427

66. Kirchdoerfer RN, Ward AB. Structure of the SARS-CoV nsp12 polymerase bound to nsp7 and nsp8 co-factors. Nat Commun. 2019 Dec;10(1):2342.

67. Decroly E, Debarnot C, Ferron F, Bouvet M, Coutard B, Imbert I, et al. Crystal Structure and Functional Analysis of the SARS-Coronavirus RNA Cap 2′-O-Methyltransferase nsp10/nsp16 Complex. Rey FA, editor. PLoS Pathog. 2011 May 26;7(5):e1002059.

68. Ma Y, Wu L, Shaw N, Gao Y, Wang J, Sun Y, et al. Structural basis and functional analysis of the SARS coronavirus nsp14–nsp10 complex. Proc Natl Acad Sci. 2015 Jul 28;112(30):9436–41.

69. Bouvet M, Lugari A, Posthuma CC, Zevenhoven JC, Bernard S, Betzi S, et al. Coronavirus Nsp10, a Critical Co-factor for Activation of Multiple Replicative Enzymes. J Biol Chem. 2014 Sep 12;289(37):25783–96.

70. Nakagawa K, Lokugamage KG, Makino S. Chapter Five - Viral and Cellular mRNA Translation in Coronavirus-Infected Cells. In: Ziebuhr J, editor. Coronaviruses [Internet]. Academic Press; 2016. p. 165–92. (Advances in Virus Research; vol. 96). Available from: http://www.sciencedirect.com/science/article/pii/S0065352716300409

71. Celniker G, Nimrod G, Ashkenazy H, Glaser F, Martz E, Mayrose I, et al. ConSurf: Using Evolutionary Data to Raise Testable Hypotheses about Protein Function. Isr J Chem. 2013;53(3–4):199–206.

72. Mayrose I, Graur D, Ben-Tal N, Pupko T. Comparison of Site-Specific Rate-Inference Methods for Protein Sequences: Empirical Bayesian Methods Are Superior. Mol Biol Evol. 2004 Sep 1;21(9):1781–91.

73. Schlessinger A, Yachdav G, Rost B. PROFbval: predict flexible and rigid residues in proteins. Bioinformatics. 2006 Apr 1;22(7):891–3.

74. Bernhofer M, Kloppmann E, Reeb J, Rost B. TMSEG: Novel prediction of transmembrane helices. Proteins Struct Funct Bioinforma. 2016;84(11):1706–16.

75. Lewis TE, Sillitoe I, Lees JG. cath-resolve-hits: a new tool that resolves domain matches suspiciously quickly. Bioinformatics. 2019 May 15;35(10):1766–7.

76. Li Y, Zhang Z, Yang L, Lian X, Xie Y, Li S, et al. The MERS-CoV Receptor DPP4 as a Candidate Binding Target of the SARS-CoV-2 Spike. iScience. 2020 Jun;23(6):101160.

## SUPPLEMENTARY REFERENCES

1. Almeida MS, Johnson MA, Herrmann T, Geralt M, Wüthrich K. Novel β-Barrel Fold in the Nuclear Magnetic Resonance Structure of the Replicase Nonstructural Protein 1 from the Severe Acute Respiratory Syndrome Coronavirus. J Virol. 2007 Apr 1;81(7):3151–61.

2. Serrano P, Johnson MA, Almeida MS, Horst R, Herrmann T, Joseph JS, et al. Nuclear Magnetic Resonance Structure of the N-Terminal Domain of Nonstructural Protein 3 from the Severe Acute Respiratory Syndrome Coronavirus. J Virol. 2007 Nov 1;81(21):12049–60.

3. Chen D, Vollmar M, Rossi MN, Phillips C, Kraehenbuehl R, Slade D, et al. Identification of Macrodomain Proteins as Novel *O* -Acetyl-ADP-ribose Deacetylases. J Biol Chem. 2011 Apr 15;286(15):13261–71.

4. Jankevicius G, Hassler M, Golia B, Rybin V, Zacharias M, Timinszky G, et al. A family of macrodomain proteins reverses cellular mono-ADP-ribosylation. Nat Struct Mol Biol. 2013 Apr;20(4):508–14.

5. Kustatscher G, Hothorn M, Pugieux C, Scheffzek K, Ladurner AG. Splicing regulates NAD metabolite binding to histone macroH2A. Nat Struct Mol Biol. 2005 Jul;12(7):624–5.

6. Forst AH, Karlberg T, Herzog N, Thorsell A-G, Gross A, Feijs KLH, et al. Recognition of Mono-ADP-Ribosylated ARTD10 Substrates by ARTD8 Macrodomains. Structure. 2013 Mar;21(3):462–75.

7. Malet H, Coutard B, Jamal S, Dutartre H, Papageorgiou N, Neuvonen M, et al. The Crystal Structures of Chikungunya and Venezuelan Equine Encephalitis Virus nsP3 Macro Domains Define a Conserved Adenosine Binding Pocket. J Virol. 2009 Jul 1;83(13):6534–45.

8. Chatterjee A, Johnson MA, Serrano P, Pedrini B, Joseph JS, Neuman BW, et al. Nuclear Magnetic Resonance Structure Shows that the Severe Acute Respiratory Syndrome Coronavirus-Unique Domain Contains a Macrodomain Fold. J Virol. 2009 Feb 15;83(4):1823–36.

9. Johnson MA, Chatterjee A, Neuman BW, Wüthrich K. SARS Coronavirus Unique Domain: Three-Domain Molecular Architecture in Solution and RNA Binding. J Mol Biol. 2010 Jul;400(4):724–42.

10. Chen Y, Savinov SN, Mielech AM, Cao T, Baker SC, Mesecar AD. X-ray Structural and Functional Studies of the Three Tandemly Linked Domains of Non-structural Protein 3 (nsp3) from Murine Hepatitis Virus Reveal Conserved Functions. J Biol Chem. 2015 Oct 16;290(42):25293–306.

11. Daczkowski CM, Dzimianski JV, Clasman JR, Goodwin O, Mesecar AD, Pegan SD. Structural Insights into the Interaction of Coronavirus Papain-Like Proteases and Interferon-Stimulated Gene Product 15 from Different Species. J Mol Biol. 2017 Jun;429(11):1661–83.

12. Bailey-Elkin BA, Knaap RCM, Johnson GG, Dalebout TJ, Ninaber DK, van Kasteren PB, et al. Crystal Structure of the Middle East Respiratory Syndrome Coronavirus (MERS-CoV) Papain-like Protease Bound to Ubiquitin Facilitates Targeted Disruption of Deubiquitinating Activity to Demonstrate Its Role in Innate Immune Suppression. J Biol Chem. 2014 Dec 12;289(50):34667–82.

13. Békés M, van der Heden van Noort GJ, Ekkebus R, Ovaa H, Huang TT, Lima CD. Recognition of Lys48-Linked Di-ubiquitin and Deubiquitinating Activities of the SARS Coronavirus Papain-like Protease. Mol Cell. 2016 May;62(4):572–85.

14. Ratia K, Kilianski A, Baez-Santos YM, Baker SC, Mesecar A. Structural Basis for the Ubiquitin-Linkage Specificity and deISGylating Activity of SARS-CoV Papain-Like Protease. Rey FA, editor. PLoS Pathog. 2014 May 22;10(5):e1004113.

15. Serrano P, Johnson MA, Chatterjee A, Neuman BW, Joseph JS, Buchmeier MJ, et al. Nuclear Magnetic Resonance Structure of the Nucleic Acid-Binding Domain of Severe Acute Respiratory Syndrome Coronavirus Nonstructural Protein 3. J Virol. 2009 Dec 15;83(24):12998–3008.

16. Xu X, Lou Z, Ma Y, Chen X, Yang Z, Tong X, et al. Crystal Structure of the C-Terminal Cytoplasmic Domain of Non-Structural Protein 4 from Mouse Hepatitis Virus A59. Masucci MG, editor. PLoS ONE. 2009 Jul 10;4(7):e6217.

17. Jin Z, Du X, Xu Y, Deng Y, Liu M, Zhao Y, et al. Structure of Mpro from SARS-CoV-2 and discovery of its inhibitors. Nature. 2020 Jun;582(7811):289–93.

18. Xue X, Yu H, Yang H, Xue F, Wu Z, Shen W, et al. Structures of Two Coronavirus Main Proteases: Implications for Substrate Binding and Antiviral Drug Design. J Virol. 2008 Mar 1;82(5):2515–27.

19. Johnson MA, Jaudzems K, Wüthrich K. NMR Structure of the SARS-CoV Nonstructural Protein 7 in Solution at pH 6.5. J Mol Biol. 2010 Oct;402(4):619–28.

20. Xiao Y, Ma Q, Restle T, Shang W, Svergun DI, Ponnusamy R, et al. Nonstructural Proteins 7 and 8 of Feline Coronavirus Form a 2:1 Heterotrimer That Exhibits Primer-Independent RNA Polymerase Activity. J Virol. 2012 Apr 15;86(8):4444–54.

21. Hillen HS, Kokic G, Farnung L, Dienemann C, Tegunov D, Cramer P. Structure of replicating SARS-CoV-2 polymerase. Nature. 2020 Aug;584(7819):154–6.

22. Kirchdoerfer RN, Ward AB. Structure of the SARS-CoV nsp12 polymerase bound to nsp7 and nsp8 co-factors. Nat Commun. 2019 Dec;10(1):2342.

23. Zhai Y, Sun F, Li X, Pang H, Xu X, Bartlam M, et al. Insights into SARS-CoV transcription and replication from the structure of the nsp7–nsp8 hexadecamer. Nat Struct Mol Biol. 2005 Nov;12(11):980–6.

24. Su D, Lou Z, Sun F, Zhai Y, Yang H, Zhang R, et al. Dodecamer Structure of Severe Acute Respiratory Syndrome Coronavirus Nonstructural Protein nsp10. J Virol. 2006 Aug 15;80(16):7902–8.

25. Joseph JS, Saikatendu KS, Subramanian V, Neuman BW, Brooun A, Griffith M, et al. Crystal Structure of Nonstructural Protein 10 from the Severe Acute Respiratory Syndrome Coronavirus Reveals a Novel Fold with Two Zinc-Binding Motifs. J Virol. 2006 Aug 15;80(16):7894–901.

26. Ma Y, Wu L, Shaw N, Gao Y, Wang J, Sun Y, et al. Structural basis and functional analysis of the SARS coronavirus nsp14–nsp10 complex. Proc Natl Acad Sci. 2015 Jul 28;112(30):9436–41.

27. Ferrer-Orta C, Sierra M, Agudo R, de la Higuera I, Arias A, Pérez-Luque R, et al. Structure of Foot-and-Mouth Disease Virus Mutant Polymerases with Reduced Sensitivity to Ribavirin. J Virol. 2010 Jun 15;84(12):6188–99.

28. Gong P, Kortus MG, Nix JC, Davis RE, Peersen OB. Structures of Coxsackievirus, Rhinovirus, and Poliovirus Polymerase Elongation Complexes Solved by Engineering RNA Mediated Crystal Contacts. Menéndez-Arias L, editor. PLoS ONE. 2013 May 8;8(5):e60272.

29. Jia Z, Yan L, Ren Z, Wu L, Wang J, Guo J, et al. Delicate structural coordination of the Severe Acute Respiratory Syndrome coronavirus Nsp13 upon ATP hydrolysis. Nucleic Acids Res. 2019 Jul 9;47(12):6538–50.

30. De I, Bessonov S, Hofele R, dos Santos K, Will CL, Urlaub H, et al. The RNA helicase Aquarius exhibits structural adaptations mediating its recruitment to spliceosomes. Nat Struct Mol Biol. 2015 Feb;22(2):138–44.

31. Zhang X, Zhan X, Yan C, Zhang W, Liu D, Lei J, et al. Structures of the human spliceosomes before and after release of the ligated exon. Cell Res. 2019 Apr;29(4):274–85.

32. Dehghani-Tafti S, Levdikov V, Antson AA, Bax B, Sanders CM. Structural and functional analysis of the nucleotide and DNA binding activities of the human PIF1 helicase. Nucleic Acids Res. 2019 Apr 8;47(6):3208–22.

33. Chakrabarti S, Jayachandran U, Bonneau F, Fiorini F, Basquin C, Domcke S, et al. Molecular Mechanisms for the RNA-Dependent ATPase Activity of Upf1 and Its Regulation by Upf2. Mol Cell. 2011 Mar;41(6):693–703.

34. Clerici M, Mourão A, Gutsche I, Gehring NH, Hentze MW, Kulozik A, et al. Unusual bipartite mode of interaction between the nonsense-mediated decay factors, UPF1 and UPF2. EMBO J. 2009 Aug 5;28(15):2293–306.

35. Lim SC, Bowler MW, Lai TF, Song H. The Ighmbp2 helicase structure reveals the molecular basis for disease-causing mutations in DMSA1. Nucleic Acids Res. 2012 Nov;40(21):11009–22.

36. Deng Z, Lehmann KC, Li X, Feng C, Wang G, Zhang Q, et al. Structural basis for the regulatory function of a complex zinc-binding domain in a replicative arterivirus helicase resembling a nonsense-mediated mRNA decay helicase. Nucleic Acids Res. 2014 Mar 1;42(5):3464–77.

37. Ferron F, Subissi L, Silveira De Morais AT, Le NTT, Sevajol M, Gluais L, et al. Structural and molecular basis of mismatch correction and ribavirin excision from coronavirus RNA. Proc Natl Acad Sci. 2018 Jan 9;115(2):E162–71.

38. Walls AC, Park Y-J, Tortorici MA, Wall A, McGuire AT, Veesler D. Structure, Function, and Antigenicity of the SARS-CoV-2 Spike Glycoprotein. Cell. 2020 Apr;181(2):281-292.e6.

39. Hakansson-McReynolds S, Jiang S, Rong L, Caffrey M. Solution Structure of the Severe Acute Respiratory Syndrome-Coronavirus Heptad Repeat 2 Domain in the Prefusion State. J Biol Chem. 2006 Apr 28;281(17):11965–71.

40. Wang N, Rosen O, Wang L, Turner HL, Stevens LJ, Corbett KS, et al. Structural Definition of a Neutralization-Sensitive Epitope on the MERS-CoV S1-NTD. Cell Rep. 2019 Sep;28(13):3395-3405.e6.

41. Yuan M, Wu NC, Zhu X, Lee C-CD, So RTY, Lv H, et al. A highly conserved cryptic epitope in the receptor binding domains of SARS-CoV-2 and SARS-CoV. Science. 2020 May 8;368(6491):630–3.

42. Xia S, Yan L, Xu W, Agrawal AS, Algaissi A, Tseng C-TK, et al. A pan-coronavirus fusion inhibitor targeting the HR1 domain of human coronavirus spike. Sci Adv. 2019 Apr;5(4):eaav4580.

43. Song W, Gui M, Wang X, Xiang Y. Cryo-EM structure of the SARS coronavirus spike glycoprotein in complex with its host cell receptor ACE2. Heise MT, editor. PLOS Pathog. 2018 Aug 13;14(8):e1007236.

44. Yan R, Zhang Y, Li Y, Xia L, Guo Y, Zhou Q. Structural basis for the recognition of SARS-CoV-2 by full-length human ACE2. Science. 2020 Mar 27;367(6485):1444–8.

45. Wang Q, Qi J, Yuan Y, Xuan Y, Han P, Wan Y, et al. Bat Origins of MERS-CoV Supported by Bat Coronavirus HKU4 Usage of Human Receptor CD26. Cell Host Microbe. 2014 Sep;16(3):328–37.

46. Li Y, Surya W, Claudine S, Torres J. Structure of a Conserved Golgi Complex-targeting Signal in Coronavirus Envelope Proteins. J Biol Chem. 2014 May 2;289(18):12535–49.

47. Surya W, Li Y, Torres J. Structural model of the SARS coronavirus E channel in LMPG micelles. Biochim Biophys Acta BBA - Biomembr. 2018 Jun;1860(6):1309–17.

48. Hänel K, Stangler T, Stoldt M, Willbold D. Solution structure of the X4 protein coded by the SARS related coronavirus reveals an immunoglobulin like fold and suggests a binding activity to integrin I domains. J Biomed Sci. 2006 May;13(3):281–93.

49. Nelson CA, Pekosz A, Lee CA, Diamond MS, Fremont DH. Structure and Intracellular Targeting of the SARS-Coronavirus Orf7a Accessory Protein. Structure. 2005 Jan;13(1):75–85.

50. Meier C, Aricescu AR, Assenberg R, Aplin RT, Gilbert RJC, Grimes JM, et al. The Crystal Structure of ORF-9b, a Lipid Binding Protein from the SARS Coronavirus. Structure. 2006 Jul;14(7):1157–65.

